# Cognitive effects of thalamostriatal degeneration are ameliorated by normalizing striatal cholinergic activity

**DOI:** 10.1101/2022.08.25.505358

**Authors:** Serena Becchi, Billy Chieng, Laura A. Bradfield, Roberto Capellán, Beatrice K. Leung, Bernard W. Balleine

**Affiliations:** School of Psychology, Faculty of Science, University of New South Wales, Sydney, Australia; School of Life Sciences, Faculty of Science, University of Technology Sydney, New South Wales, Australia; School of Psychology, Department of Psychobiology, National University for Distance Learning, Madrid, Spain

**Keywords:** Parafascicular thalamus, dorsomedial striatum, goal-directed action, neuroinflammation, degeneration, cholinergic interneurons, ATPase activity

## Abstract

The loss of neurons in parafascicular thalamus (Pf) and of their inputs to dorsomedial striatum (DMS) are associated with Lewy body disease (LBD) and Parkinson’s disease dementia (PDD) and have been linked to the effects of neuroinflammation. In rats, these inputs regulate the function of striatal cholinergic interneurons (CINs) that are necessary for the flexible encoding of the action-outcome (AO) associations for goal-directed action. We found that these inputs modify the encoding, not retrieval, of new AO associations and cause burst-pause firing of CINs in the DMS during AO remapping. These adaptive effects were abolished by neuroinflammation in the Pf, resulting in the loss of goal-directed control when the rats were required to update AO associations after a change in contingency. We found that the neuronal and behavioral deficits induced by inflammation in the Pf were rescued by administration of selegiline, a MAO-B inhibitor that we found also enhances ATPase activity in CINs, suggesting a potential treatment for cognitive deficits associated with inflammation affecting the function of midline thalamic nuclei and related structures.

## INTRODUCTION

The ability to encode, retrieve and update knowledge of the consequences of our actions is necessary for goal-directed actions to remain adaptive in complex environments. Learning new strategies to achieve valued goals requires the capacity to encode specific action-outcome (AO) relationships, the accurate retrieval of which is necessary to capitalize on that information when the appropriate circumstances arise. Nevertheless, environments can change, sometimes temporarily, sometimes more permanently, and so flexibility of encoding is necessary to update previous strategies in these situations without necessarily erasing previously learned ones.

There is now considerable evidence that the dorsomedial striatum serves to coordinate flexibility in the encoding and retrieval of the specific AO associations for goal-directed action in these situations *(1, 2)*. When initially encountered, the encoding of these relations appears to depend on plasticity at direct pathway striatonigral spiny projection neurons *(3)*, driven by activity in the cortico-striatal pathway *(4)* and dependent on the modulatory influence of local dopamine release in this area *(5)*. However, recent evidence suggests that remapping this learning as the environment changes depends on inputs to the dorsomedial striatum (DMS) from the parafascicular thalamic nucleus (Pf) via the thalamostriatal pathway, which functions to control the interdigitation of new learning with previously learned strategies in a manner that reduces interference between them *(6, 7)*.

The primary targets of the thalamostriatal pathway are the giant aspiny cholinergic interneurons (CINs) *(8–10)* that comprise less than 3% of striatal neurons but, through their broad arborization and mosaic distribution, generate the highest level of acetylcholine in the brain *(11, 12)*. Pf-control of the burst-pause firing pattern of CINs in the DMS *(13)* is vital for altering both cholinergic and dopaminergic function within targeted territories of the striatum, for encouraging or discouraging plasticity in those territories and so for encoding new, and protecting pre-existing, AO information when conditions in the environment change *(6, 7)*. Thus, damage to the Pf-DMS pathway has been found to render existing and new learning subject to mutual interference leaving animals incapable of appropriately engaging in goal-directed action control. Given the functional importance of this pathway, its degeneration in dementias associated with Parkinson’s disease (PD) and dementia with Lewy bodies (LBD), is of considerable concern *(14)*. In PD the progressive loss of dopaminergic neurons in the nigrostriatal pathway is accompanied by up to 30-40% of neuronal loss in Pf area *(15–18)*. This latter neurodegeneration appears to be unrelated to motor deficits and contributes more to cognitive dysfunctions in LBD *(19)* and both the early, premotor stages, and advanced stages of PD when patients develop Parkinson’s disease dementia (PDD) *(20–22)*. Similarly, in models of PD, monkeys chronically treated with 1-methyl-4-phenyl-1,2,3,6-tetrahydropyridine (MPTP) show intralaminar thalamic degeneration and a decrease in their terminals in the striatum *(19)*. However, preclinical studies on the neuropathological basis of PDD and LBD are rare – particularly with regard to effects on the thalamostriatal pathway – mainly due to a lack of suitable animal models for dementia-related cognitive dysfunction. Thus, the question of how to model such dysfunction in animals remains critical.

The aetiology of the influence of PDD and LBD on the thalamostriatal pathway is unclear, but a role for chronic inflammation has been widely proposed as a possible mechanism *(23)*. Neuroinflammation is increasingly considered a contributor to dementia pathogenesis *(24)*, and is associated with the accumulation of Lewy body deposits, which are also particularly evident in the Pf area *(18)*. Moreover, direct evidence of inflammation in LBD is growing, with elevated microglial activation identified at post-mortem *(25)* and *in vivo* using positron emission tomography (PET) scan *(26)*.

Here, we built upon this accumulating evidence to conduct a series of experiments using a disease-relevant model of cognitive dysfunction in rats that directly relies on dysfunction of the thalamostriatal pathway, induced by targeted injections of the pro-inflammatory agent lipopolysaccharide (LPS) into the Pf. Given the complex dopaminergic and cholinergic profile of CIN-related modulation of striatal principal neurons, we investigated the effects of selegiline, a monoamine-oxidase inhibitor, with a known cholinergic profile, originally used to palliate initial motor symptoms of early stages of PD, on the deficits in neuronal and cognitive function induced by inflammation in the Pf. We report here that the administration of selegiline during AO remapping does indeed rescue both the neuronal, learning- and memory-related deficits in goal-directed action control, normalizing CIN activity and goal-directed control after inflammation-induced damage to the Pf and Pf-DMS pathway.

## RESULTS

### The Pf-DMS pathway is necessary to encode but not to retrieve changes in the AO association for goal-directed action

Considerable anatomical evidence has confirmed the existence of a direct Pf-posterior DMS (pDMS) pathway that projects unilaterally but extensively throughout all portions of the dorsal striatum *(6, 27–29)*. Despite converging evidence that this pathway mediates the updating of AO associations *(6)*, direct causal evidence is still lacking. In particular, it is not currently known whether this pathway mediates the updating of learning or the retrieval of updated associations after learning. Therefore, to establish the precise role of the Pf-pDMS pathway in AO-updating for our subsequent studies in this series, we first addressed this question using a chemogenetic approach. We stereotaxically injected a retro-Cre virus in the pDMS to express Cre recombinase in pDMS-projecting Pf neurons (**Fig. 1A**). We separated rats into four groups: in three groups a Cre-inducible hM4 DREADD virus was then stereotaxically injected into the Pf to promote the viral-mediated expression of hM4D, a Gi/o-coupled muscarinic M4 DREADD (**Fig. 1A-C**). This receptor is activated by the intraperitoneal (ip) injection of an artificial ligand clozapine-N-oxide (CNO) to induce membrane hyperpolarization and neuronal silencing *(30)* in transfected Pf neurons projecting to pDMS, as demonstrated by electrophysiological recording (**Fig. 1D, E**). A fourth group received an infusion of a null virus to serve as a CNO control.

**Fig. 1.**
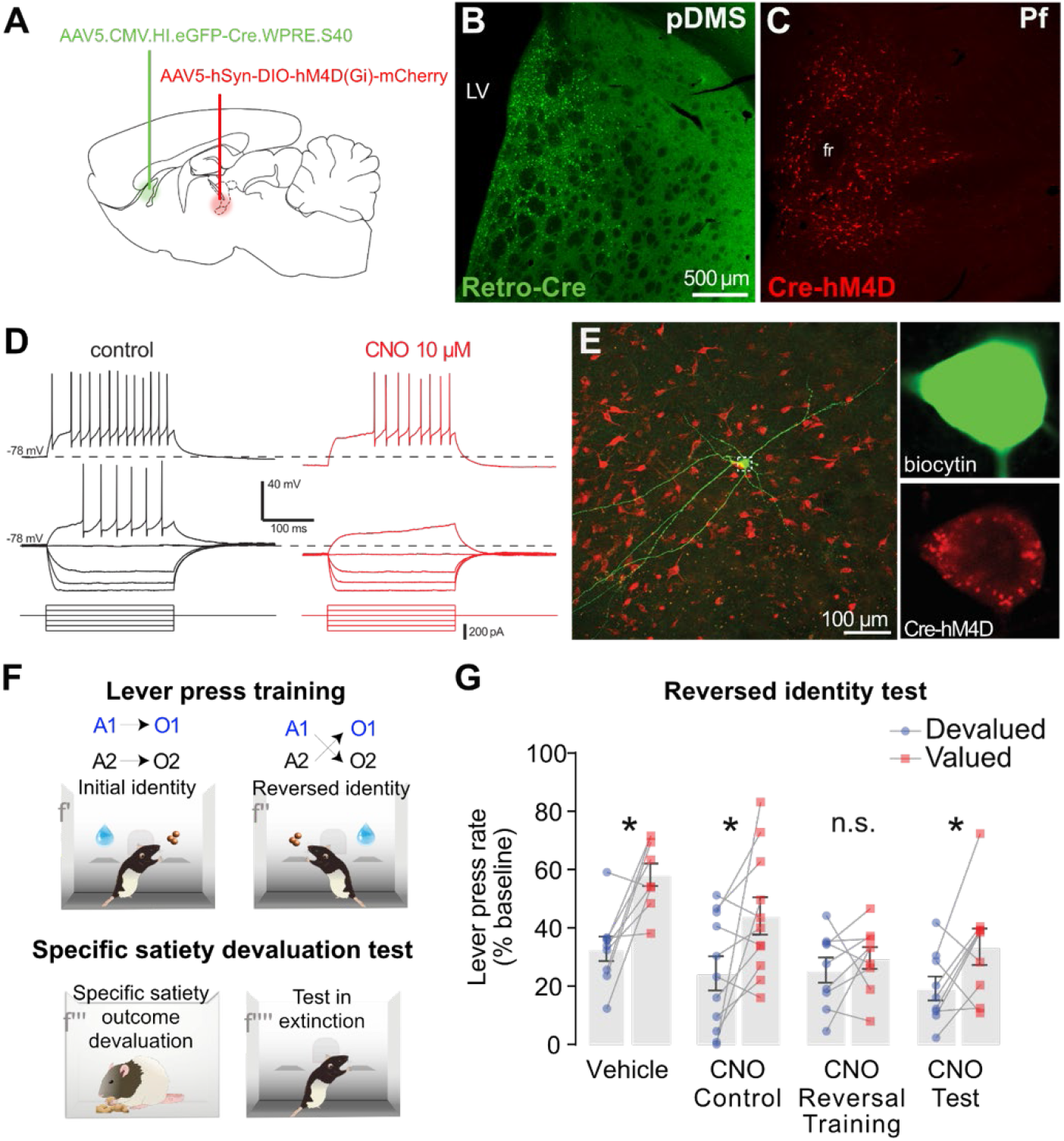
The role of the Pf-DMS pathway is encoding and retrieving changes in the AO contingency. **(A)** Illustration of viral infusions given in Pf (red) and pDMS (green). **(B)** Representative micrographs of the injection site and the spread of Retro-Cre in pDMS, and **(C)** of Cre-hM4D in the Pf. **(D)** Confirmation that CNO reduced the excitability of Pf neurons expressing hM4Di (n=3). Under whole-cell current clamp, cell was resting at −78mV before drug superfusion. A family of voltage-current relationships was then sampled by injection of incremental step current pulses. Traces were in the same cell before (left, black) and during (right, red) application of CNO (10μM). Voltage recording from the most depolarized step is separated for clarity. **(E)** Micrograph of a biocytin-filled thalamostriatal Pf neuron from the recordings shown in (D). **(F)** Design of the behavioral procedures used in this and subsequent experiments: After LPS injections in the Pf, rats were trained on two action-outcome (AO) associations (Initial identity) (f’). They were then trained on contingencies for which the outcomes associated with the two actions were reversed (Reversed identity – f”). Rats were then given specific satiety-induced outcome devaluation (f’”) followed by a choice test on the two levers conducted in extinction (f””). **(G)** Outcome devaluation test after reversal training to assess the flexibility of goal-directed actions in rats expressing hM4D that received vehicle injections (Vehicle n=9), rats with mCherry and CNO injections (CNO Control n=11), rats with hM4D and CNO injections during reversal training sessions (CNO Reversal Training n=9) and rats with hM4D and CNO injections at test (CNO test n=9). Graphs represent means ±1 SEM; *indicates p<0.05.

Rats were then given behavioral training – **Fig. 1F** (described in Bradfield et al, 2013 *(6)* – see Methods) with rats in groups Vehicle (M4+Vehicle), CNO Control (mCherry+CNO), CNO Reversal Training (M4+CNO-Reversal Training), and CNO Test (M4+CNO-Test) trained to press one lever for grain pellets and a second lever for a 20% sucrose solution (counterbalanced) with reward delivery on increasing ratio schedules. All groups acquired lever pressing similarly: **Fig. S1A – initial identity**; there was an effect of training, F(3,34)=64.41, p<0.001, that did not interact with group, F<1. They were then trained on the reverse of these contingencies; i.e., if initially the left lever earned pellets and the right lever sucrose then the left lever now earned sucrose and the right lever pellets. Rats in CNO Reversal Training and CNO Control groups received ip injections of CNO (7mg/kg) 1 hr prior to reversal lever press sessions whereas rats in CNO Test and Vehicle groups received saline injections at the same time. Responding during the reversal stage is shown in **Fig. S1B - identity reversed**. Again reversal training did not differ across groups: there was an effect of training, F(2.517, 34)=30.83, p<0.001, that did not interact with group, F(7.552, 34)=1.43, p>0.05 (Greenhouse-Geisser corrected for violations of sphericity).

To evaluate what was learned after identity reversal, rats received an outcome devaluation test for which they were pre-fed one of the outcomes (pellets or sucrose) for 1 hr to induce sensory specific satiety (**Fig. 1F**)*(2)*. Immediately prior to being placed in the devaluation feeding boxes, rats in groups CNO Test and CNO Control received ip injections of CNO whereas rats in groups CNO Reversal Training and Vehicle received ip injections of saline. Following specific satiety induction, rats were transferred to the operant boxes for a 5min devaluation extinction test in which both levers were extended but neither earned any outcomes. The test data are shown in **Fig. 1G**. From this figure, it is clear that devaluation was intact (i.e., valued > devalued) for both control groups, as well as group CNO Test, but was impaired (valued = devalued) for group CNO Reversal Training. To analyze these data (see methods) we first compared the control groups (Vehicle and CNO Control) and then the controls vs. CNO Test. This analysis revealed a main effect of devaluation, F(1,34)=21.26, p<0.001, but no interaction between either the Vehicle and CNO Control groups or when comparing these controls with the CNO Test group (Fs<1). In contrast, the comparison of these three groups with the CNO Reversal Training group revealed a devaluation x group interaction, F(1,34)=4.65, p=0.038. Follow up simple effects analyses of this interaction confirmed that devaluation was intact (valued > devalued) for groups Vehicle, F(1,34)=12.98, p=0.001, CNO Control, F(1,34)=9.51, p=0.004 and CNO Test, F(1,34)=4.083 p=0.051, but not for group CNO Reversal Training, F<1.

Together, these results suggest that the direct Pf→pDMS pathway is required for learning to interlace new and existing AO contingencies, but, once this has been achieved, appears not to be required for the retrieval of this information.

### Effect of neuroinflammation in the Pf on cholinergic activity in the pDMS

We have explored several means of causing targeted disruption of the Pf→pDMS pathway in animal models, from excitotoxic lesions of the Pf in the thalamus of rats *(6)* to chemogenetic silencing described above. In the following experiments, we assessed whether we could induce a comparable effect on this pathway using neuroinflammation. Our first model used targeted injections of the pro-inflammatory agent lipopolysaccharide (LPS), which, when delivered in the Pf, caused a clear degeneration of the thalamostriatal pathway. This was revealed by retrograde tracing using Cholera Toxin B (CTB) (**Fig. S2A-C**). Immunohistological analysis revealed a decrease of CTB-positive neurons, indicating that LPS-induced neuronal loss included pDMS projecting neurons in the Pf. Direct LPS infusion in the Pf caused microglia activation and recruitment along with neuronal loss (**Fig. 2A-D**). Quantification of NeuN particles in the Pf showed that LPS induced a moderate but significant loss of NeuN-positive neurons, p=0.006, unpaired *t*-test (t=3.056, df=20; **Fig. 2E**).

**Fig. 2.**
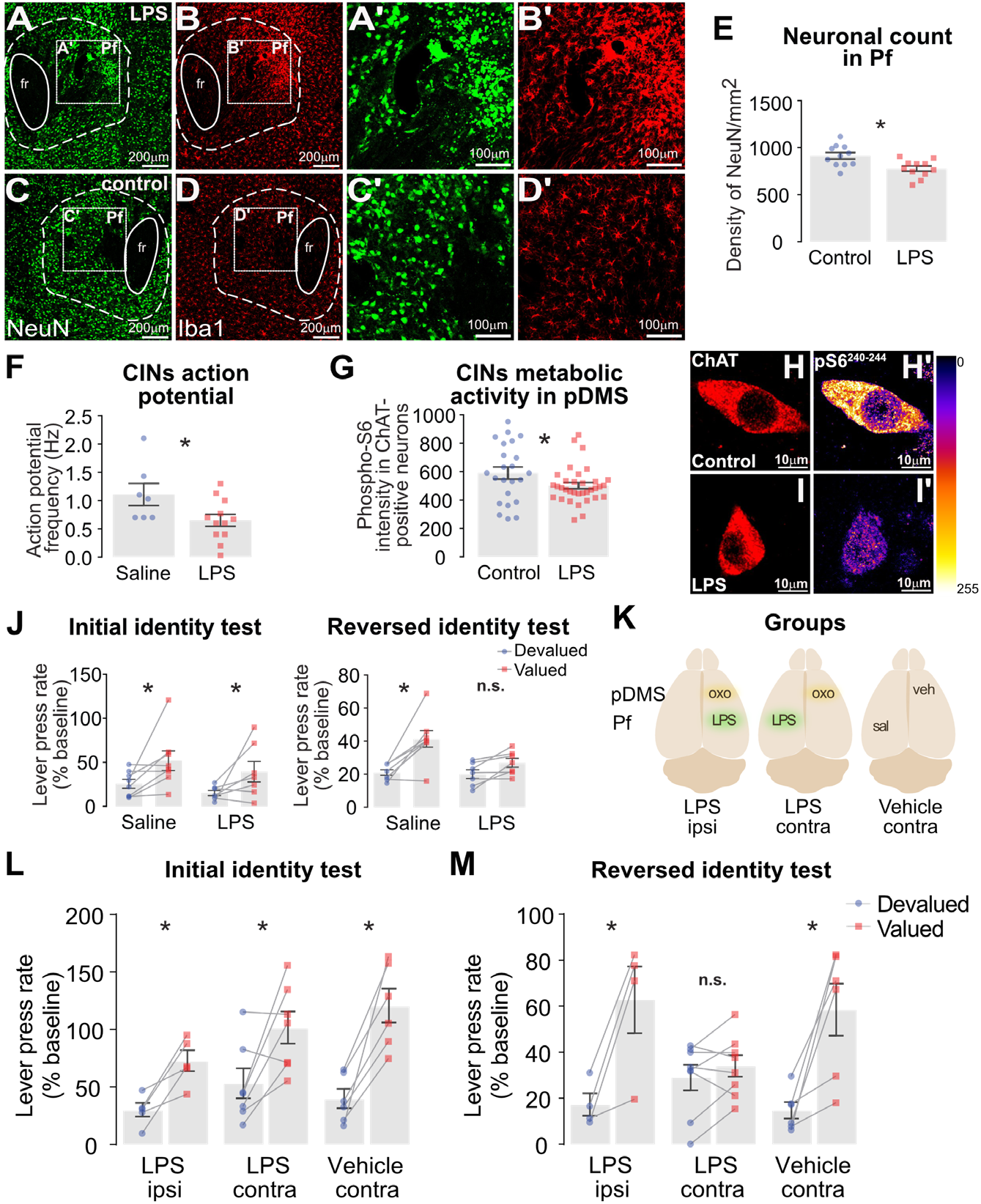
Neuroinflammation in the Pf and its effect on CIN activity in the pDMS and on behavioral function. **(A-D)** Representative micrographs of LPS infusion (A,B) and control hemisphere (C,D) in the Pf on the left and insets shown on the right **(A’-D’)**. Sections were labelled for microglia (Iba1-red) and neurons (NeuN-green). **(E)** Quantification of neuronal density in the Pf. Control n=13, LPS n=13. **(F)** Spontaneous action potential frequency in CINs sampled under cell-attached recording conditions from *ex vivo* brain slices taken from saline and LPS pretreated rats. Saline n=7, LPS n=12. **(G)** The metabolic activity of CINs calculated via quantification of p-S6rp protein activity at residue 240-244. Saline n=23 cells from 7 rats, LPS n=34 cells from 12 rats **(H-I)** Representative micrographs of CINs, identified by ChAT-positive staining, and p-S6rp protein expression. **(H’-I’)** A pseudocolor palette (Lookup Table, LUT) highlights the intensity of p-S6rp fluorescence (display range: 0–255). **(J)** Outcome devaluation test after initial AO identity training (left) and after the AO identity was reversed (right), in rats that received either Saline or LPS injected in the Pf. Lever press rate is expressed as a percentage of lever pressing during training (% baseline). Saline n=8, LPS n=7. **(K)** Illustration of disconnection experimental design: Rats received Oxo-S in the pDMS and LPS in the Pf either ipsilaterally (LPS ipsi) or contralaterally (LPS contra), the former to retain and the latter to disconnect the Pf-pDMS (CIN) pathway during reversed contingency learning. A third control group received saline in the Pf and vehicle in the pDMS injected contralaterally (Vehicle contra). **(L)** Lever press rate, expressed as % baseline, during a choice test conducted after initial identity learning, and **(M)** after the outcome identities were reversed, in these same three groups. Vehicle contra n=6, LPS ipsi n=6, LPS contra n=9. Error bars reflect ±1 SEM; * p<0.05.

Next, we examined the effect of LPS in the Pf on the function of CINs in the pDMS. For this experiment, rats were given bilateral injections of LPS or saline. After 2 weeks we performed patch-clamp *ex vivo* electrophysiology on 300μm coronal slices centered on the pDMS. Sampling CINs in the rat pDMS based on electrophysiological characteristics, morphology and *post hoc* immunohistochemistry, we found that action potential frequency was significantly reduced in CINs recorded from the LPS infused hemisphere compared with CINs recorded from the saline infused hemisphere, p=0.038, unpaired two-tailed *t*-test (t=2.257, df=17; **Fig. 2F**). To confirm that the inflammation-induced reduction in firing rate was specific to changes in the intrinsic activity of CINs, we measured functionally relevant changes in CIN activity based on fluctuations in phosphorylation levels of the ribosomal protein S6 (p-S6rp) assessed using immunofluorescence *(31)* (**Fig. 2G-I**). We quantified the state of phosphorylation of C-terminal 240-244 residues of ribosomal protein S6, an integrant of the ribosomal complex modulated in striatal CINs. In the hemisphere with LPS injection in the Pf, we detected a reduction in the p-S6rp signal in CINs in the pDMS ipsilateral to the inflammation compared to control hemispheres, p=0.049, unpaired two-tailed *t*-test (t=2.013, df=55; **Fig. 2G**). Together, these results suggest that glutamatergic neurons in the Pf-pDMS pathway are sensitive to Pf inflammation resulting in the impairment of CIN activity in the pDMS.

### LPS-induced inflammation in the Pf impairs AO updating after a shift in contingency

We next examined the effect of bilateral infusion of LPS into the Pf on goal directed learning after a shift in the AO contingency using outcome identity reversal. We used a similar behavioral protocol to that described above – **Fig. 1F** (ref to Bradfield et al., 2013) *(6)*. Rats given bilateral LPS or saline infusions into the Pf were food deprived and trained on a two lever-two outcomes procedure. Both groups learned to press the levers and increased performance with the increasing response requirement over the course of training (**Fig. S2D**). Importantly, in this study we first assessed the encoding of the initial outcome identities using an outcome devaluation assessment before progressing to identity reversal. We found that LPS had no effect on initial AO encoding: (**Fig. 2J - Initial identity test**): Contrast analysis revealed an overall effect of devaluation F(1,13)=9.843, p=0.008 but no main effect of group F(1,13)<1 and no group x devaluation interaction F(1,13)<1.

We then assessed the ability of the rats to update their goal-directed learning and show flexibility in AO encoding by reversing the identity of the outcomes earned by the two levers (**Fig. S2D - Reversed**). Despite the LPS group showing higher pressing rates, both groups earned equal number of rewards during reversal training (**Fig. S2E**). To assess what the rats learned during identity reversal, we gave them a second outcome devaluation test at the end of reversal training and found, in marked contrast to the test after the initial learning, the LPS group failed to show an outcome devaluation effect (**Fig. 2J – Reversed identity test)**. Again contrast analysis supported these observations, revealing a significant group × devaluation interaction, F(1,13)=6.307, p=0.026 with simple effect analyses finding a reliable difference between devalued and valued levers in the saline group, F(1,13)=30.808, p<0.001, but not in the LPS group, F(1,13)=3.074, p>0.05. Thus, as found with NMDA Pf lesions *(6, 32)* LPS-induced neuroinflammation in the Pf impairs goal-directed action only after a change in the AO contingency.

### The role of CINs in the effect of neuroinflammation in the Pf on AO remapping

To establish whether the impairment in interlacing new with existing AO associations observed in the LPS group depended on an inflammation-induced effect on cholinergic activity in the pDMS, we disconnected the Pf-pDMS pathway by infusing LPS unilaterally into the Pf prior to initial training and infused oxotremorine (Oxo-S) contralaterally into the pDMS during identity reversal training. Oxo-S is a selective muscarinic agonist that binds to the muscarinic M2/M4 autoreceptors expressed on CINs *(6)*, inhibiting the functions of these cells *(6, 33–35)*. For this experiment, we compared a group that received LPS injected in the Pf and Oxo-S in the contralateral pDMS (LPS contra) with two control groups: one that received these infusions ipsilaterally (LPS ipsi) and a saline control (Vehicle contra) that receiving saline infusions contralaterally (see **Fig. 2K**). We conducted a direct replication of the behavioral procedures described above and all groups responded similarly during initial acquisition (**Fig. S2F)** and showed evidence that they learned the initial AO associations, as revealed by an outcome devaluation test of initial identity encoding (**Fig. 2L**). Contrast analysis revealed a main effect of devaluation, F(1,18)=43.386, p<0.001, but no devaluation x group interaction was found when comparing the two control groups (Vehicle contra vs. LPS ipsi), F(1,18)=3.938, p=0.063 or control groups (Vehicle contra+ LPS ipsi) vs. LPS Contra, F(1,18)=1.876, p=0.188.

We then trained the rats for four sessions with outcome identities reversed. Prior to each session of training on the new contingencies, rats were given an infusion of either Oxo-S or vehicle. As has been reported previously *(6)*, Oxo-S produced a mild non-selective reduction in performance in both LPS ipsi and LPS contra infused groups (**Fig. S2F**). After reversal training, rats received a drug free outcome devaluation test. Contralateral infusions of Oxo-S during training produced a clear deficit in encoding the new AO contingency: rats that received these infusions in pDMS contralaterally to the LPS infusion in Pf pressed both levers at a similar rate on test, whereas rats given intra-pDMS infusions of Oxo-S ipsilaterally to Pf-LPS or of vehicle rats showed a reliable outcome devaluation effect (valued > devalued; **Fig. 2M**). Contrast analysis revealed a main effect of devaluation F(1, 18)=47.09, p<0.001, and no interaction when comparing the two control groups (Vehicle contra vs. LPS ipsi) x devaluation F<1. However, when the control groups combined were compared against LPS contra, this analysis revealed a significant interaction with devaluation, F(1, 18)=18.32, p<0.001. Simple effect analysis showed clear preference for the valued lever over the devalued lever in Vehicle contra rats, F(1, 18)=29.049 p<0.001 and LPS ipsi rats, F(1,18)=26.66 p<0.001, whereas performance did not differ significantly in the LPS contra rats, F(1,18)<1, p>0.05.

### Reversed contingency training increases burst-pause firing activity in CINs

Having established the importance of CINs and their function as a target of the thalamostriatal pathway, we next sought to establish how the activity of CINs is affected by activity in the Pf-pDMS pathway and, subsequently, by both initial identity training and by reversed identity training. First, we infused channelrhodopsin into the Pf and stimulated Pf terminals in the pDMS while under whole-cell recording of a CIN. Optical illumination of the brain slice (LED 473 nm, 4 mW, 20 Hz, pulse duration 5 ms) modified the regular firing of spontaneous action potentials to elicit a burst-pause firing pattern – **Fig. 3A-B.**

**Fig. 3.**
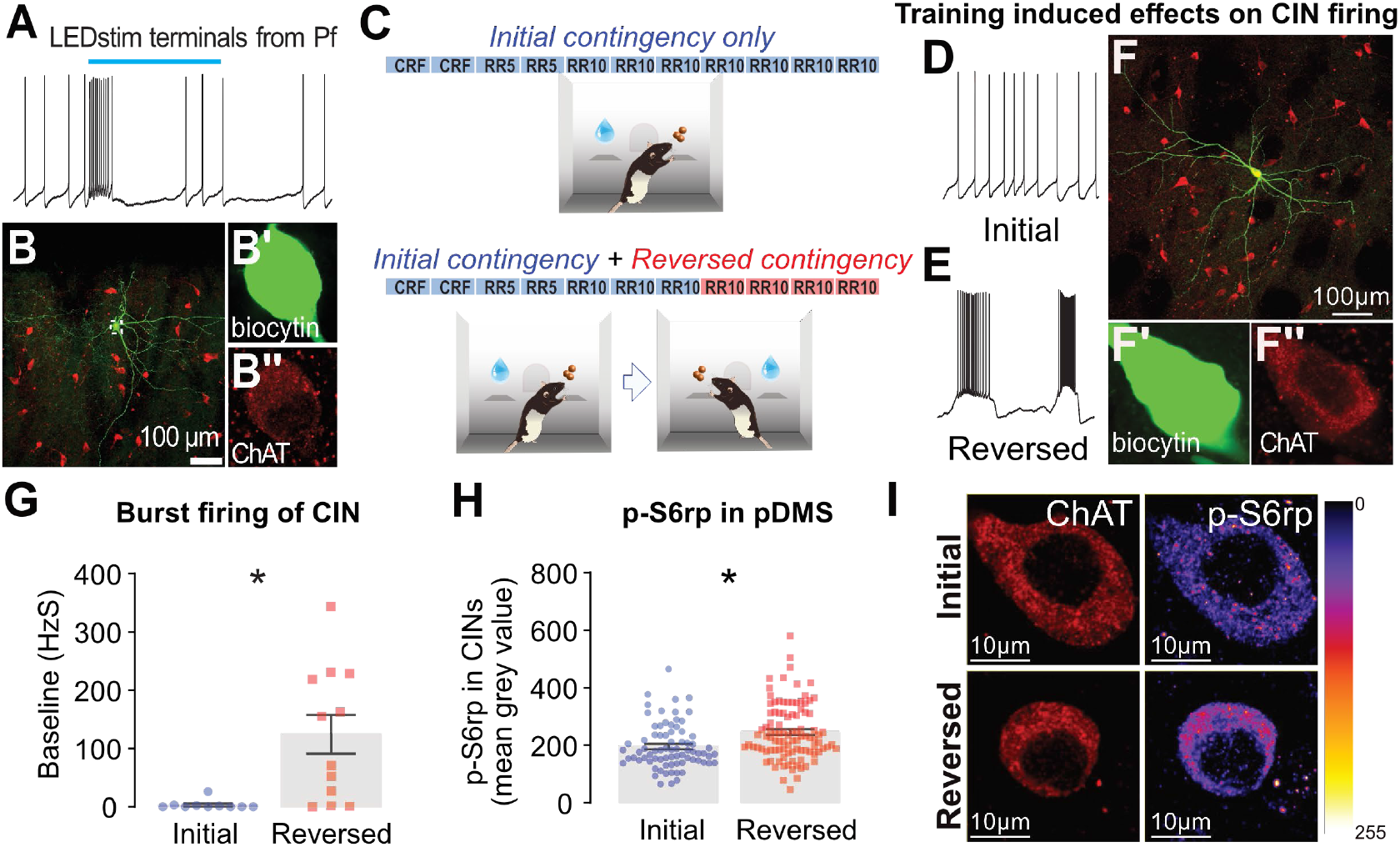
Stimulation of the Pf-pDMS pathway and training on reversed AO contingencies induces burst-pause firing activity in CINs in the pDMS. **(A-B)** Stimulation of thalamostriatal terminals in pDMS evokes burst-pause firing in CINs. Under whole-cell recording in a CIN, optical illumination of the brain slice (LED 473nm, 4mW, 20Hz, pulse duration 5ms) elicited a burst-pause firing pattern after quite regular firing of spontaneous action potentials. Blue bar on top denotes duration (5s) of LED light applied (A). Below is the micrograph for the corresponding CIN recorded (B), typically labelled with biocytin (green) after the recording shown in (A) and *post hoc* histology revealing ChAT positive immunoreactivity (red). (B’) and (B”) are enlargements of the cell body in (B). **(C)** Experimental design: Half of the rats were trained on the initial AO association (Top) whereas the other half were trained similarly and then given outcome identity reversal (Bottom). **(D-E)** Representative voltage recording traces showing spontaneous action potential firing in CINs from rats trained under the initial (D) and reversed (E) contingencies. **(F)** A typical CIN labelled with biocytin (green) during whole-cell recording and later *post hoc* histology revealing ChAT positive immunoreactivity (red). (F’) and (F”) are enlargements of the cell body in (F). **(G)** ‘Burst firing score’ aggregates (for analysis see Methods) quantified from spontaneous action potential measurements under whole-cell recording of CINs from rats trained under the initial and reversed contingencies. Initial n=10 cells from 8 rats, Reversed n=12 cells from 10 rats. **(H)** CIN metabolic activity via quantification of p-S6rp. Initial n=76 cells from 10 rats, reversed n=100 cells from 10 rats. **(I)** Representative micrographs of high-magnification confocal images showing p-Ser240-244-S6rp intensity in ChAT-immunoreactive neurons in the pDMS. LUT highlights the intensity of p-S6rp fluorescence (display range: 0–255). Error bars reflect ±1 SEM. * denotes p<0.05.

Next, we gave two groups of rats the same amount of training except one group was maintained on the initial AO contingencies throughout whereas the second group was given four days exposure to outcome identity reversal – **Fig. 3C (Figure S3** for training data). Immediately after the final session of instrumental training, brain slices were taken for *ex vivo* electrophysiology. The electrophysiology was performed by an experimenter blind to the group from which the slices has been taken. Standard whole-cell patch clamp assessment of CINs in the pDMS found a significant increase in the burst-pause pattern of action potential firing in CINs in the group given outcome identity reversal compared to rats for whom the initial contingency was maintained (unpaired *t*-test t=3.325, df=20, p=0.003; **Fig. 3D-G**). In a separate cohort, we also found significantly higher p-S6rp immunoreactivity in CINs after reversed contingency trained rats (unpaired *t*-test t=3.727, df=174, p<0.001; **Fig. 3H-I**). We have previously reported that p-S6rp activity at residue 240-244 reflects changes in CIN firing, with increased activity associated with burst firing *(31)*. Together, therefore, these findings suggest that the functional engagement of CINs, as reflected in their burst-pause activity, shows the engagement of the Pf-pDMS pathway in the remapping/updating of AO encoding and the interlacing of that new learning with previous learning *(36, 37)*.

### The MAO-B inhibitor selegiline increases burst-pause activity in CINs

We next sought to understand how changes in Pf-pDMS pathway modify the burst-pause activity of CINs. We focused on the role of dopamine, which plays a central role in most aspects of cellular activity within the striatum and particularly in striatal cholinergic signaling and CIN excitability. Action potential firing in CINs can lead to terminal release of dopamine in the striatum via nicotinic cholinergic receptors *(38)*. On the other hand, CINs have dopaminergic D1 and D2 receptors, as demonstrated by responses from selective receptor agonists and antagonists *(6, 39–41)*. Elevating synaptic dopamine using cocaine has been reported to enhance the pause in CIN action potential firing generating a more pronounced burst-pause pattern, whereas the D2 receptor antagonist sulpiride shortens the pause *(37)*. Based on these findings, we first sought to explore the relationship of dopamine to burst-pause firing in CINs by comparing the effect of the monoamine oxidase (MAO)-B inhibitor selegiline (100μM) with that of the MAO-A/B inhibitor pargyline (100μM) on CIN activity in *ex vivo* brain slices from untrained naïve rats. Selegiline, but not pargyline, significantly increased the burst-pause firing pattern in CINs, with some neurons demonstrating a more dramatic effect than others (non-parametric Friedman test shows Friedman chi-square=12.67, df=8, p=0.001; **Fig. 4A**). Follow up Dunn’s multiple comparison revealed a difference between baseline and selegiline (p=0.001), but not between baseline and pargyline (p>0.05).

**Fig. 4.**
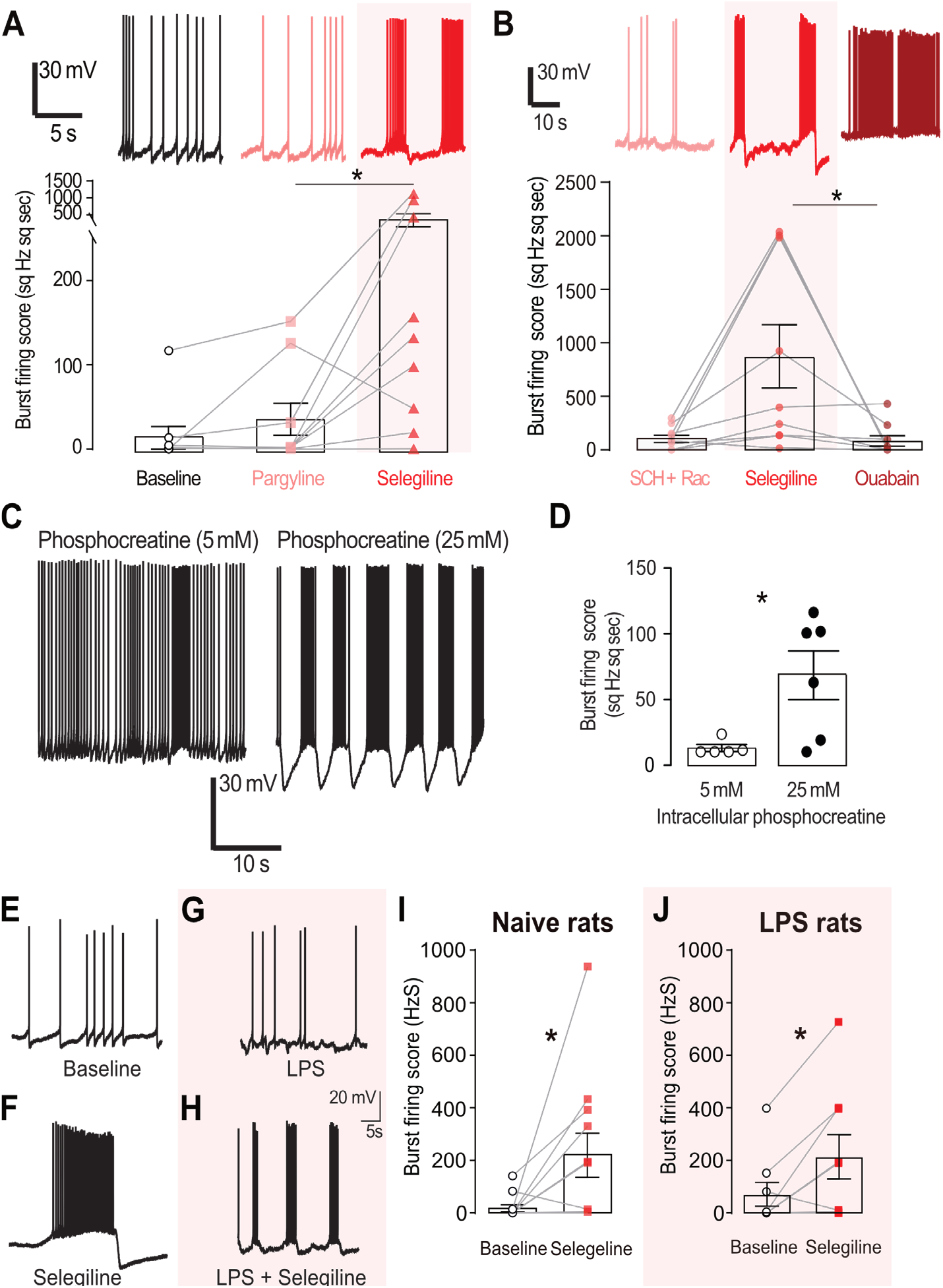
The MAO-B inhibitor Selegiline enhances burst-pause firing in CINs. **(A)** Selegiline (100μM), and not pargyline (100μM), significantly increased burst-pause firing in CINs (n=9). **(B)** The selegiline-induced (100μM) increase in burst-pause firing persisted in the presence of SCH23390 (SCH, 10μM) and raclopride (Rac, 10μM), but was abolished by ouabain (100μM, n=9). **(C)** Raw traces showing that a higher concentration of phosphocreatine (25mM) in the recording pipette significantly increased burst-pause firing in CINs. **(D)** Grouped data for 5 mM (n=5) and 25mM (n=6) phosphocreatine in the intracellular solution. **(E-F)** Selegiline increased burst-pause firing in CINs in untrained naive rats (n=11) and **(G-H)** in rats with LPS injected into the Pf (n=9). Panels **(E-H)** show raw traces and **(I-J)** show grouped data. Error bars represent ±1 SEM. * denotes p<0.05.

To varying degrees, there is often some burst-pause and irregular firing in CINs at baseline *(42, 43)*. However, we found that the selegiline-induced burst-pause firing pattern in CINs persisted in the presence of SCH23390 (10μM, D1 antagonist) and raclopride (10μM, D2 antagonist) but was blocked by the sodium-potassium ATPase blocker ouabain (100μM, **Fig. 4B**) showing a reliable group effect (Friedman chi-square=8.667, df=8, p=0.010). Dunn’s multiple comparisons revealed a trend in the difference between D1/D2 antagonist and selegiline (p=0.101), a significant difference between ouabain and selegiline treatments (p=0.014) but no difference between ouabain and D1/D2 antagonists (p>0.05) suggesting the burst-pause action potential firing evoked by selegiline was not dopamine-mediated and appeared instead to be related to a metabolic mechanism.

We have previously reported that elevated firing in CINs can increase p-S6rp immunoreactivity, a metabolic process that involves mTOR *(31)*. Activity in CINs is also intricately linked to the availability of ATP, with these neurons shutting down completely when the level of ATP is lowered by opening ATP-sensitive potassium channels *(44)*. The routine practice of incorporating a small amount of phosphocreatine (5-10mM) in the intracellular pipette solution is thought to regenerate ATP in viable brain slice experiments *(45)*. In the present study, we found that a higher concentration of phosphocreatine (25mM) significantly increased the burst-pause firing pattern in CINs (Unpaired *t*-test, t=2.685, df=9, p=0.025; **Fig. 4C-D**). In rat mesencephalic trigeminal neurons, sodium-potassium ATPase has been reported to interact with the hyperpolarization-activated cation current (Ih)*(46)*, but we did not find any evidence that Ih in CINs was affected by selegiline application (p>0.05; **Fig. S4**). Nevertheless, selegiline’s ability to increase the burst-pause firing pattern in CINs persisted in both naïve (**Fig. 4E-F,I**) and in rats given an infusion of LPS into the Pf (**Fig. 4G-H,J**). Statistical analysis revealed that selegiline increased burst firing in both the naïve rats (paired *t*-test, t=2.369, df=10, p=0.0394) and LPS-infused rats (paired *t*-test, t=2.605, df=8, p=0.031).

### LPS-induced impairment in AO contingency updating is rescued by selegiline

Based on the finding that selegiline increases burst-pause firing in CINs, we hypothesized that its administration could rescue the behavioral impairment produced by neuroinflammation in the Pf induced by LPS infusion. To assess this hypothesis, we conducted a series of experiments in which we either injected selegiline systemically or infused it bilaterally into the pDMS during outcome-identity reversal training.

In delivering selegiline peripherally, rats received bilateral injections of LPS or saline into the Pf as described previously – **Fig. 5A**. After 2 weeks, rats were given instrumental training for the initial AO identity wherein LPS rats showed similar acquisition and sensitivity to outcome devaluation to saline controls (**Fig. S5A, 5B**): Analysis of the devaluation test showed a main effect of devaluation F(1,43)=43.098, p<0.001 but no group x devaluation interaction F<1. Next, rats were trained with the outcome identities reversed. Prior to each daily session, they received an injection of either selegiline (1mg/kg) or saline ip, creating four groups. Finally, we assessed whether the rats were able to update the AO contingencies using a second outcome devaluation test. As is clear from **Fig. 5C**, whereas the devaluation effect in the saline (Sal Sal) and saline+selegiline (Sal SEL) controls did not differ, it was again abolished in the LPS Sal group. However, this debilitating effect of LPS was not observed in rats given an injection of selegiline prior to identity reversal training (LPS SEL) which showed a clear devaluation effect. Statistical analysis found an effect of devaluation, F(1,42)=27.87, p<0.001, but no interaction between the controls (Sal Sal vs. Sal SEL) or between the controls combined and the LPS SEL group, (Fs<1). In contrast, comparison of these three groups (Sal Sal + Sal SEL + LPS SEL) against LPS-Sal found a group x devaluation interaction, F(1, 42)=4.803, p=0.034. Follow up simple effect analysis between devalued and valued levers revealed a significant difference in the Sal Sal group, F(1,42)=13.218, p=0.001, the Sal SEL group F(1,42)=9.562, p=0.004 and the LPS SEL group F(1, 42)=9.185, p=0.004 but not the LPS Sal group, F<1. To confirm the effect of ip selegiline on CIN function in the pDMS, we examined p-S6rp intensity in ChAT+ neurons in the pDMS of rats from the four groups immediately before a fifth training session and found an increase of p-S6rp signaling after selegiline treatment (**Fig. S5B**).

**Fig. 5.**
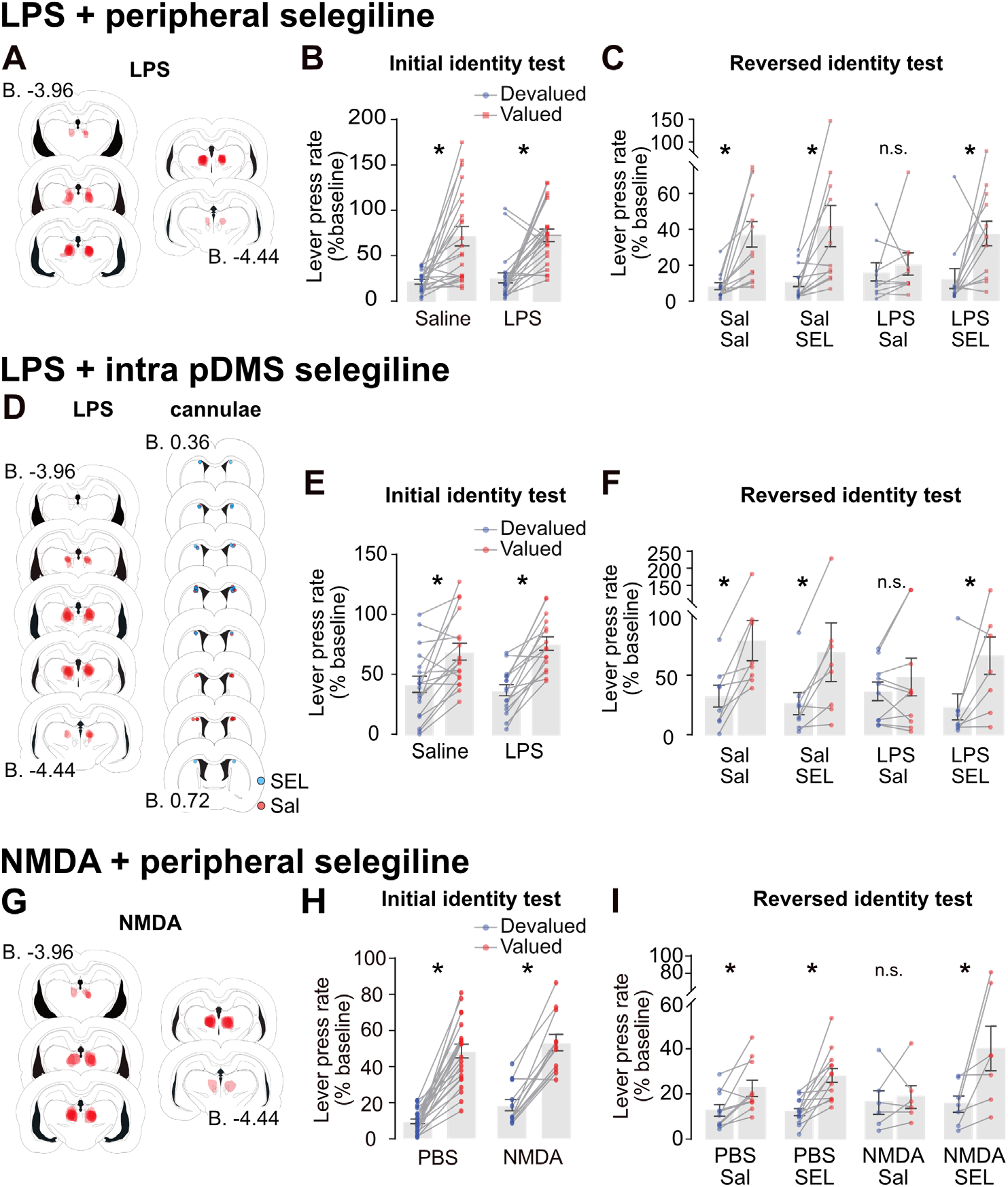
Action-outcome contingency update impairment can be rescued by administration of Selegiline. **(A)** Reconstruction of LPS lesion in the Pf. **(B**) Lever press rate during outcome devaluation expressed as percentage to responding during training (% baseline) following initial AO identity in rats that received bilateral injections of either saline or LPS in the Pf and **(C)** after the reversed identity training, where saline (Sal Sal, LPS Sal) or selegiline (Sal SEL, LPS SEL) was administered systemically prior to each reversal training session. Sal Sal n=13, Sal SEL n=11, LPS Sal n=12, LPS SEL n=10. **(D)** Maps of LPS infusions in the Pf and cannula placements in the pDMS. **(E)** Lever press rate during outcome devaluation as percentage baseline following initial AO identity training in rats that received bilateral injections of either saline or LPS in the Pf and **(F)** after the reversed identity training where saline (Sal Sal, LPS Sal) or selegiline (Sal SEL, LPS SEL) was infused in the pDMS prior to each reversal training session. Sal Sal n=8, Sal SEL n=8, LPS sal n=10, LPS SEL n=8. **(G)** Spread of NMDA lesion in the Pf. **(H)** Lever press rate during outcome devaluation as percentage baseline following initial AO identity training in rats that received bilateral injections of either NMDA or PBS in the Pf and **(I)** after the reversed identity training where saline (PBS Sal, NMDA Sal) or selegiline (PBS SEL, NMDA SEL) was administered systemically prior to each reversal training session. PBS Sal n=10, PBS SEL n=13, NMDA Sal n=6, NMDA SEL n=7. Error bars represent ±1 SEM. * denotes p<0.05.

We next sought to establish whether the effect of peripherally administered selegiline could be replicated through microinfusions directly into the pDMS (see methods). Accordingly, all groups and methods were the same as the previous experiment except selegiline or saline was locally infused into the pDMS 20 min prior to each outcome identity reversal training session. LPS affected areas and cannula placements in the pDMS are shown in **Fig. 5D**. There were no differences in training on the initial and reversed contingencies between groups that received LPS or saline in the Pf (**Fig. S5B**). In the first devaluation test, LPS again had no effect on goal-directed behavior: contrast analysis revealed an overall devaluation effect F(1, 30)=51.30 p<0.001 but neither an effect of group or an group x devaluation interaction, F(1,30)=1.65, p>0.05 (**Fig. 5E**).

Importantly, similar to a peripheral selegiline injection, we again found that selegiline was able to rescue the LPS-induced deficit in AO encoding during identity reversal. Rats given microinfusions of selegiline during reversal training performed similarly to controls (**Fig. 5F**). Contrast analysis again supported these observations: there was an overall devaluation effect F(1,30)=28.22, p<0.001. We found the size of the devaluation effect in the two saline groups (Sal Sal vs Sal SEL) did not differ (Fs<1) and nor did it differ these controls were compared with the LPS group that received Selegiline (LPS SEL; F<1). However, the devaluation effect in these three groups combined (Sal Sal + Sal SEL + LPS SEL) differed significantly from the LPS Sal group, F(1, 30)=4.716, p=0.038. Comparison between devalued and valued levers revealed a reliable simple effects in the Sal Sal group, F(1,30)=11.125, p=0.002, the Sal SEL group F(1,30)=9.523, p=0.004, the LPS SEL group F(1,30)=9.483, p=0.004, but not in the LPS Sal group, F<1, p>0.05.

Finally, we sought to extend these findings and assess whether selegiline was able to reverse the behavioral impairment in encoding flexibility previously observed in another model of thalamic degeneration induced by the infusion of a cytotoxic concentration of NMDA into the Pf *(6)*. Rats received bilateral injections of NMDA or PBS into the Pf (**Fig. 5G**). In a subset of rats that received NMDA, we first assessed the effect of selegiline on CIN firing and found that incubation of selegiline with NMDA lesioned brain slices induced burst firing activity (**Fig. S5D**). Next, we trained and tested the remaining rats on the same behavioral protocol as described in the previous experiments (**Fig. 5H-I, S5E**). NMDA lesions had no effect on initial acquisition of goal-directed behavior (**Fig. 5H**), generating a main effect of devaluation F(1, 34)=158.9, but neither an effect of group F(1,34)=3.324, p>0.05 nor a group x devaluation interaction F<1. In the final devaluation test after reversal learning, however, we found a deficit in the NMDA lesioned rats that received saline injections during reversal training (NMDA Sal – **Fig. 5I**) that was, again, ameliorated in rats that received injections of selegiline during reversal training (NMDA SEL). Contrast analysis revealed an overall devaluation effect F(1, 32)=24.813, p<0.001, and a main effect of selegiline F(1, 32)= 6.962, p=0.013, but no difference in the size of the devaluation effects in the control groups (PBS Sal vs. PBS SEL) or when these groups were combined and compared to the NMDA SEL group, Fs<1. However, the three groups (PBS Sal + PBS SEL + NMDA SEL) compared to NMDA Sal generated a group x devaluation interaction F(1, 32)=4.406, p=0.044. Comparison between devalued and valued levers revealed a reliable simple effect for the PBS Sal group, F(1,32)=4.307, p=0.046, the PBS SEL group F(1,32)=14.281, p<0.001, the NMDA SEL group, F(1,32)=18.125, p<0.001, but no difference for the NMDA Sal group, F(1,31)<1.

## DISCUSSION

For goal-directed actions to remain adaptive, specific AO relationships have to be learned and encoded in such a manner that they can be individually retrieved without interfering with each other. Any such interference would be highly maladaptive: it would render the consequences of actions uncertain, hinder decision-making, block or at least make tenuous action selection particularly when contingencies become uncertain in a changing context. The current experiments explored the role of the thalamostriatal pathway in AO encoding in these changing circumstances using an outcome-identity reversal task designed to discriminate failures of AO learning from effects on attention or novelty/familiarity. Importantly, using this task, we found this pathway was necessary to remap or update current AO learning but was not necessary for performance after such encoding confirming the role of this pathway in allowing new AO associations to be encoded without interference from those already encoded. These findings suggest that neurological conditions resulting in degeneration of this pathway will have significant effects on goal-directed action control. We evaluated this prediction using neuroinflammation and NMDA-induced cytotoxicity models of degeneration and found direct evidence that degeneration of this pathway abolished goal-directed control but only when animals were forced to update previous learning. The projection from the thalamus likely has many effects, but we found its influence on AO encoding depends predominantly on its influence on cholinergic activity in the pDMS, particularly on the firing activity of CINs. Finally, we found that the influence of this degeneration could be ameliorated by correcting its effects on CIN firing, providing a treatment for the degeneration-induced cognitive deficits resulting in the maladaptive control of goal-directed action.

More specifically, we first established, using an hM4 DREADD expressed on pDMS projecting neurons in the Pf, that activity in the Pf–pDMS pathway is necessary during the encoding of new AO associations and not during their retrieval. We then investigated the influence of neuroinflammation in the Pf, induced by the infusion of LPS, on AO encoding. We found that LPS caused considerable changes in Pf and degeneration of the thalamostriatal pathway, similar to that we found in the case of dopamine neurons when infused into the substantia nigra *(47)*. However, it did not affect initial AO encoding and only abolished the ability of rats to encode AO associations after AO associations changed during outcome identity reversal, as we found previously after NMDA lesions of the Pf *(6, 7)*. At a cellular level, activity in the thalamostriatal projection has been found to induce a burst-pause firing pattern in CINs (35, 36, 46) and we were able to replicate this effect using ChR2 infused into the Pf and stimulating Pf inputs to the pDMS. Guided by this finding we then sought to decouple the Pf input and its effects on CIN activity using an asymmetrical disconnection approach. We infused LPS unilaterally into the Pf in one hemisphere and inhibited striatal CIN activity in the contralateral striatum using the cholinergic M2/4 autoreceptor agonist oxotremorine. Relative to controls, disconnection had a similar functional effect to bilateral LPS and blocked AO encoding after outcome identity reversal.

Although the mechanics of burst firing are intrinsic to CINs *(42, 43)*, an important role for the thalamostriatal input may be to initiate this burst-pause activity under specific conditions or after changes in state *(32, 49)*, such as that induced by a shift in AO contingency. Consistent with this hypothesis, we found that, in an *ex vivo* slice preparation, CINs exhibited significantly more burst-pause firing immediately after reversed-contingency training than in rats maintained under the original contingency. Concurrently, p-S6rp immunoreactivity at residue 240-244 in CINs was elevated in the reversed-contingency group. Conversely, CINs recorded in slices taken from rats with LPS infused into the Pf showed reduced p-S6rp immunoreactivity and decreased spontaneous firing. Thus, activity in the thalamostriatal pathway appeared to be sufficient to provide the necessary excitatory drive to induce striatal CINs to fire in a burst-pause pattern and so to modulate plasticity during reversed-contingency learning.

The burst-pause action potential firing in CINs is known to be at least partially regulated by dopamine *(37)* and we evaluated its role by comparing the effects of the selective MAO-B inhibitor selegiline with that of the MAO A/B inhibitor pargyline on CIN firing. We found that selegiline, but not pargyline, could induce burst-pause firing in CINs from *ex vivo* striatal slices taken from untrained naïve rats. However, surprisingly, this effect appeared not to be dependent on dopamine activity: selegiline was found to continue to induce burst-pause activity in the presence of dopamine D1 and D2 antagonists. Furthermore, we found that selegiline was able to normalize burst pause firing in CINs on slices taken from rats given LPS infusions into the Pf. Selegiline does have other pharmacological actions, although less well reported. Perhaps most importantly, it is an enhancer of sodium potassium ATPase activity, an energy-demanding enzyme that consumes a large amount of ATP to maintain electrogenicity and excitability in mammalian cells *(50)*. This suggests that selegiline may induce its effects by increasing intracellular ATP to stimulate sodium potassium ATPase activity and ultimately burst firing in CINs. In support of this hypothesis, we found that ouabain, an inhibitor of sodium potassium ATPase, abolished Selegiline-induced burst firing. By contrast, elevating intracellular ATP level via incorporation of the ATP regenerative agent phosphocreatine inside the recording pipette significantly increased burst firing in CINs. Unfortunately, ouabain affects all mammalian cells, and we have so far been unable to induce a change in sodium potassium ATPase that is limited to CINs. Burst-pause firing in CINs involves activation of many calcium channels and calcium stores *(51)*. Whilst blocking calcium entry and depleting stores can inhibit burst-pause firing in CINs from *ex vivo* brain slice experiments, these drugs can also directly affect activity of spiny projection neurons. Lastly, the spontaneous firing of CINs is critically dependent on normal functioning of a number of ion channels and the cationic channel Ih has been attributed to the spontaneous depolarization phase needed by CINs for their unique spontaneous activity *(52, 53)*. We found selegiline had no effect on Ih in CINs.

Together these data suggested that selegiline may be able to reverse the functional effects of LPS-induced degeneration of the thalamostriatal pathway on goal-directed action, specifically on AO encoding after changes in the action-outcome contingency. To assess this, we first examined the effect of peripherally administered selegiline in rats given either LPS-induced or NMDA-induced degeneration of the Pf. We found that injections of selegiline during new learning, when outcome identities were reversed, rescued the ability of rats to accurately encode these reversed contingencies as assessed in a subsequent outcome devaluation test. Importantly, we found a similar effect when selegiline was administered centrally directly into the pDMS, confirming the locus of selegiline’s effect.

Selegiline (aka l-deprenyl) has been used clinically as an adjunct to treat early-stage PD in an attempt to delay the introduction of l-dopa for control of motor movement. Its effect on the cognitive effects of Parkinson’s disease or other dementias is more equivocal with some studies reporting an improvement and others ambiguous results. This is understandable given the to dementia disease variants recruited in these studies, mostly with unknown neurological degeneration characteristics. Whilst our investigations demonstrated a significant improvement in encoding new action-outcome contingencies it will be important to extend these findings to other capacities. In accord with our attempt to establish the specific function of the thalamostriatal pathway and its role in controlling striatal activity, our experiments applied manipulations to healthy CINs, as validated by electrophysiological recordings from test subjects, with only one brain region inactivated at a given time. In human drug trials there is often multi-focal neuronal degeneration in the diseased brain, and this could dilute the effectiveness of drugs, such as selegiline, in overcoming deficits, producing mixed results. It will be important therefore to investigate the effects of selegiline in broader disease models. Nevertheless, the results of the current study, in which selegiline was able to completely reverse the effects of a deficit in a complex cognitive learning process mediating goal-directed action, appear promising and suggest a potential drug treatment for neuroinflammation-induced cognitive decline.

## MATERIALS AND METHODS

### Study design

The objective of this study was to explore the effect of inflammation in the Pf on the Pf-pDMS pathway and the encoding of new AO associations for goal-directed action. We first used a chemogenetic approach to inactivate the Pf-pDMS pathway during new AO learning. Subsequently, we used LPS-induced inflammation in the Pf to induce inactivation and a disconnection approach to confirm this inflammation affected striatal cholinergic activity. Next, we assessed the importance of CINs in the pDMS for the acquisition of new AO associations and, using *ex vivo* electrophysiology, we examined the effect of new AO learning on CINs burst-firing. In a series of *ex vivo* studies we then assessed whether selegiline affects CIN burst-firing and the cellular basis for this effect before assessing whether either ip or intra-striatal selegiline could rescue the behavioral deficits induced by bilateral LPS and NMDA-induced degeneration of the Pf. Some animals were excluded from the analysis based on histological assessment. The final number of samples is indicated in the figure legends. In electrophysiological and immunohistochemical analyses investigators were blind to experimental group.

### Study approval

All procedures were approved by the University of New South Wales Ethics Committee. All animal studies reported are in compliance with the ARRIVE guidelines *(54, 55)*.

### Animals

Females and males Long-Evans rats, weighing between 250-350g females and 400-500g at the beginning of the experiment, were used as subjects. Rats that experienced behavioral training and testing were maintained at ~85% of their free-feeding body weight by restricting their food intake to between 8 and 12g of their maintenance diet per day.

### Surgeries

Rats were anaesthetized with isofluorane (5% for induction and 2–3% for maintenance) and positioned in a stereotaxic frame (Kopf, Model 942). An incision was made to expose the scalp and the incisor bar was adjusted to align bregma and lambda on the same horizontal plane. For all rats, holes were drilled into the skull above the appropriate targeted structures using coordinates in millimeters and relative to bregma and skull. Before the start of the surgery the animals received injections of an antibiotic (Benacillin, 0.3-0.4ml sc) and a local anesthetic (Bupivicaine, 0.1ml sc) at the surgical site. During surgery, rats received an injection of an analgesic (Carprofen 0.03-0.04ml sc) and warm saline (5ml ip).

For behavioral experiments, microinfusions were delivered using a microinjector (Harvard apparatus, 11 Elite) using a 1μl or 5μl Hamilton syringe at an infusion rate of 0.1 μl/min. After each injection, the needle was left in situ for an additional 6 min to avoid reflux along the injection track. In the pDMS (−0.1 anteroposterior (AP), ±2.5 mediolateral (ML), −4.7 dorsoventral (DV))*(56)*, rats received injections of AAV5.CMV.HI.eGFP-Cre.WPRE.SV40 (Addgene #105545, 0.3μl). In the Pf (females: −4.1 AP; ±1.3 ML; −6.1mm DV; males: −4.4 AP; ±1.3 ML; −6.55 DV)*(56)*, rats received injections of LPS (Sigma #L2880; 5mg/ml in sterile Saline, 1.2ul), NMDA (Sigma #M3262; 10mg/mL in sterile PBS 0.1M, 0.5μl) or AAV5-hSyn-DIO-hM4D-mCherry (Addgene #44362, 0.75μl). Rats in control groups received either rAAV5/hSyn-DIO-mCherry (Addgene #50459, 0.75μl), saline (1.2μl) or sterile PBS (0.5μl).

For retrograde tracing, rats received microinjections of Cholera Toxin B (CTB) in the pDMS (-0.5 AP, +2.5 ML, −4.7 DV)*(56)*. CTB was delivered via a glass micropipette attached to the end of a nanoject (Drummond Scientific company, Nanoject II). Six 18.4nL of CTB (List labs #104) were injected at 30s intervals. The micropipette was left in place for a further 10min to allow the CTB to diffuse.

For intracranial infusions of Oxo-S and selegiline in the pDMS, rats received a 26-gauge guide cannula (P1Technologies #C315GRL/SP) implanted above the pDMS. The tip of the guide cannula was aimed at 1 mm above the target region (i.e. −0.0 AP, ±2.5ML, −3.5 DV)*(56)*. The guide cannulas were maintained in position with screws and dental cement, and dummy cannulas were kept in each guide at all times except during microinfusions. At the time of infusions, 33G infusion cannulas (P1Technologies #C315I/SP) were lowered and Oxo-S (Sigma #O9126, 1μM, 1μL), selegiline (Sigma, #M003, 1mM, 1μL) or saline (1μl) was infused using a syringe pump (Harvard Apparatus, PHD Ultra) at a rate of 0.4μl/min with 2 min diffusion before being removed.

### Behavioral Procedures

#### Training and Devaluation

##### Magazine Training

On day 1, all rats were placed in operant chambers for approximately 20 min. In each session of each experiment, the house light was illuminated at the start of the session and turned off when the session was terminated. No levers were extended during magazine training. 20 grain pellets and 20 20% sucrose solution outcomes were delivered to the magazine port on an independent random time (RT) 60s schedule.

##### Lever Training

The animals were next trained to lever press on random ratio schedules of reinforcement. Each lever was trained separately each day and the specific lever-outcome assignments were fully counterbalanced. The session was terminated after 20 outcomes were earned or after 30 min. For the first 2 days, lever pressing was continuously reinforced. Rats were shifted to a random ratio (RR)-5 schedule for the next 2 days (i.e., each action delivered an outcome with a probability of 0.2), then to an RR-10 schedule (or a probability of 0.1) for 3 days. Devaluation Extinction Tests. After the final day of RR-10 training, rats were given free access to either the pellets or the sucrose solution for 45-min in the devaluation cage. The aim of this pre-feeding procedure was to satiate the animal specifically on the pre-fed outcome, thereby reducing its value relative to the non-prefed outcome *(2)*. Rats were then placed in the operant chamber for a 5 min choice extinction test. During this test, both levers were extended and lever presses recorded, but no outcomes were delivered. The next day, a second devaluation test was administered with the opposite outcome. Rats were then placed back into the operant chambers for a second 5 min choice extinction test.

##### Contingency Reversal Training

Subsequent to the extinction test, rats were trained to lever press on an RR-10 schedule with the previously trained contingencies reversed. That is, the lever that previously earned grain pellets now earned sucrose solution, and the lever that previously earned sucrose solution now earned pellets. Contingency reversal training continued for 4 days.

### Immunofluorescence

Rats were anaesthetized with sodium pentobarbital (150 mg/kg ip, Virbac Pty. Ltd., Australia) and transcardially perfused with 400mL of 4% paraformaldehyde (PFA) in sodium phosphate buffer (0.1M PB; pH 7.4). Brains were post-fixed overnight in 4% PFA PB and stored at 4°C. Coronal sections (35μm) were cut with a vibratome (Leica Microsystems VT1000) and stored at −20°C in a solution containing 30% ethylene glycol, 30% glycerol and PB, until they were processed for immunofluorescence.

Free-floating sections were rinsed in Tris-buffered saline with sodium fluoride (TBS-NaF; 0.25 M Tris, 0.5 M NaCl and 0.1mM NaF, pH 7.5), incubated for 5 min in TBS-NaF containing 3% H2O2 and 10% methanol, and then rinsed 10 min three times in TBS-NaF. After 20 min incubation in 0.2% Triton X-100 in TBS-NaF, sections were rinsed three times in TBS-NaF again. Choline acetyltransferase (ChAT) and the double phosphorylated form of S6 ribosomal protein (p-Ser240-244 -S6rp) were simultaneously detected through incubation with combined polyclonal goat anti-ChAT (1:500, #AB144P, Millipore) and polyclonal rabbit anti-p-Ser240-244 -S6rp (1:500, #2215, Cell Signaling Technology) primary antibodies diluted in TBS-NaF (4°C, overnight). NeuN and Iba-1 were simultaneously detected through incubation with combined polyclonal mouse anti-NeuN (1:1000, #MAB377, Merk) and polyclonal goat anti-Iba-1 (1:1000, #AB5076, Abcam) primary antibodies diluted in TBS (4°C, overnight). Sections were then rinsed 10 min in TBS three times and incubated 60 min with donkey anti-goat Alexa-594 (#A11058, Invitrogen) and donkey anti-mouse Alexa-488 (#A21202, Invitrogen) secondary antibodies diluted 1:1000 in TBS. Sections were rinsed four times 10 min in TBS before being mounted on slides and coverslipped with Vectashield fluorescence medium (H-1000-10, Vector Laboratories).

Another set of Pf sections with LPS injection were double stained with NeuN and CTB through incubation with combined polyclonal mouse anti-NeuN (1:1000, #MAB377, Merk) and polyclonal goat anti-CTB (1:1000, #703, Listlabs) primary antibodies diluted in TBS (4°C, overnight). Sections were then rinsed 10 min in TBS three times and incubated 60 min with donkey anti-goat Alexa-594 (#A11058, Invitrogen) and donkey anti-mouse Alexa-488 (#A21202, Invitrogen) secondary antibodies diluted 1:1000 in TBS. Sections were rinsed four times 10 min in TBS before being mounted on slides and coverslipped with Vectashield fluorescence medium (H-1000-10, Vector laboratories).

Fluorescence analysis and cell counts images were obtained using sequential laser scanning confocal microscopy (Fluoview FV1000, BX61WI microscope, Olympus). All ChAT-immunoreactive neurons in dorsomedial and dorsolateral striatal regions were detected in the microscope using a 60X objective (UPFL 60X oil) and centered in the acquisition area. Focal plane with optimal ChAT immunoreactivity was determined in channel 2 (Ch02; HeNe green laser). Sequential high-resolution images (optical magnification: 60X; digital zoom: 4X; resolution: 1024×1024 px) were obtained for p-Ser240-244 -S6rp signal (Ch01) and corresponding ChAT signal (Ch02) with a Kaplan filter (5 averaging scans). Laser intensity, PMT voltage and offset were maintained constant in all acquisitions of the same experiment. Raw 16-bit images were then analyzed using ImageJ software (MacBiophotonics upgrade v. 1.43u, Wayne Rasband, National Institutes of Health, USA). In superimposed channels, somatic neuronal area (excluding the nucleus) was determined as ROI in Ch02 (ChAT signal) and mean fluorescence intensity (mean grey value, grey values ranging from 0 to 255) was measured in corresponding Ch01 (p-Ser240-244 -S6rp signal). A pseudo-color palette highlighting intensity of fluorescence (16-color LookUp Table, display range: 0-255) was applied to representative neurons.

NeuN-positive neurons in Pf region were quantified as follows: high-resolution scanning confocal images of Pf were obtained from unilateral LPS injected rats or control rats, imaging the same coronal level (optical magnification: 10X; resolution: 2048×2048 px). NeuN-positive cell counts were performed using imageJ software (Cell counter plugin). Prior to all quantifications, all image files in each experiment were randomly renumbered using a MS Excel plug-in (Bio-excel2007 by Romain Bouju, France). An ROI of the Pf was drawn to exclude the needle track in LPS injected sides and the image was converted to 8-bit. A thresholding was calculated, then, after applying a Gaussian blur filter (sigma radius 1), the image was made binary and the watershed plug-in was applied. The cell counter plug-in calculated the neurons that were in the size range of 50-400μm^2^ and circularity of 0.20-1.00. Iba-1 staining was used to show the effect of LPS and to assess correct placements of injections.

### Brain slice preparation

Rats were euthanized under deep anesthesia (isoflurane 4% in air), and their brains were rapidly removed and cut on a vibratome in ice-cold oxygenated sucrose buffer containing (in mM): 241 sucrose, 28 NaHCO3, 11 glucose, 1.4 NaH2PO4, 3.3 KCl, 0.2 CaCl2, 7 MgCl2. Coronal brain slices (300μm thick) containing the pDMS or Pf were sampled and maintained at 33°C in a submerged chamber containing physiological saline with composition (in mM): 130 NaCl, 2.6 KCl, 1.4 NaH2PO4, 1.2 MgCl2, 2.5 CaCl2, 12 glucose and 27 NaHCO3, and equilibrated with 95% O2 and 5% CO2.

### Electrophysiology

After equilibrating for 1h, slices were transferred to a recording chamber and visualized under an upright microscope (Olympus BX50WI) using differential interference contrast (DIC) Dodt tube optics, and superfused continuously (1.5 ml min-1) with oxygenated physiological saline at 33°C. Cell-attached and whole-cell patch-clamp recordings were made using electrodes (2–5 MOhm) containing internal solution consisting of the following (in mM): 115 K gluconate, 20 NaCl, 1 MgCl2, 10 HEPES, 11EGTA, 5 phosphocreatine di(tris) salt (#P1937, Sigma-Aldrich), 5 Mg-ATP, and 0.33 Na-GTP, pH7.3, osmolarity 285–290mOsm l-1. Biocytin (0.1%, #B4261, Sigma-Aldrich) was added to the internal solution for marking the sampled neurons during whole-cell recording. Data acquisition was performed with a Multiclamp 700B amplifier (Molecular Devices), connected to a Macintosh computer and interface ITC-18 (Instrutech). Action potential deflections in cell-attached configuration and voltage/current recordings under whole-cell configuration were sampled at 5kHz (low pass filter 2kHz; Axograph X, Molecular Devices). Data from cell-attached and whole-cell recordings were only included in analyses if (1) the neurons appeared healthy under DIC on the monitor screen, (2) cholinergic interneurons were spontaneously active during cell-attached recording, (3) action potential amplitudes were at least 60 mV above threshold after establishing whole-cell recording mode, and (4) neurons demonstrated physiological characteristics of cholinergic interneurons such as the presence of hyperpolarization-activated cation current Ih but no plateau low-threshold spiking *(52)*, to ensure that only highly viable neurons were included. Liquid junction potentials of -10 mV were not corrected. All current-clamp data were recorded at resting membrane potential.

### Action potential analysis

To calculate an aggregate as ‘burst firing score’ for each neuron, in Axograph X, variance of spike (or action potential) instantaneous frequency is multiplied by variance of inter-spike interval. Baseline was taken as just before drug application (analysis of 200 spikes), whilst drug effect was typically taken around 5-10 min mark during application (analysis of 200 spikes). For a CIN that undergoes intense burst firing, there will be a large variance in spike frequency and also a large variance in inter-spike interval. In ouabain, drug effect was taken after 1.5 min because action potentials completely disappeared after 5 min application.

### Post hoc histology

Immediately after physiological recording, brain slices containing biocytin-filled neurons were fixed overnight in 4% PFA/0.16 M PB solution and then placed in 0.5% Triton X-100/PB for 3 d to permeabilize cells. Slices were then placed in 10% horse serum/PB for 1 h before being incubated in primary goat anti-ChAT (1:500; #AB144P, Merck) for 2 d at 4°C to aid identification of CINs. The slices were rinsed in PB and then in a one-step incubation containing both Alexa Fluor 647-conjugated donkey anti-goat secondary antibody (1:500; #A21447, Invitrogen) and Alexa Fluor 488-conjugated Streptavidin (1:1000; #S11223, Invitrogen) for 2 h. For slices containing Pf neuron plus mCherry labelling, after cell permeation with Triton X-100, they were incubated only in Alexa Fluor 488-conjugated Streptavidin (1:1000; #S11223, Invitrogen) for 2h. Stained slices were rinsed in PB 3 times for 10 min each, mounted/dried on glass slide, and coverslipped with Fluoromount-G mounting medium (#0100-01 Southern Biotech). Neurons were imaged under a confocal microscope (Fluoview FV1000 and BX61WI, Olympus).

### Drugs

All the following drugs, except CNO, were purchased from Sigma-Aldrich, St Louis, MO. Working drug concentrations: LPS (O55:B5, L2880, Sigma, 5mg/ml dissolved in PBS, 1 μl), oxotremorine sesquifumarate (Oxo-S, #O9126, Sigma, 1μM dissolved in PBS), selegiline (R(-)-deprenyl HCl, #M003, Sigma, 1mg/kg ip and 1mM, 1μL infused in the pDMS), clozapine-N-oxide (CNO, RTI International, custom synthesis batch# 13626-76) 7 mg/kg ip. Drugs for electrophysiology, selegiline (100μM), R(+)-SCH23390 HCl (D054, 10μM), S(-)-raclopride (+)-tartrate (#R121, Sigma, 10μM), ouabain octahydrate (#O3125, Sigma, 100μM), pargyline HCl (#P8013, Sigma, 100μM), and CNO (#13626-76, RTI International, 10μM).

### Statistical analysis

Experiments with two groups were analysed using two-tailed Student’s *t*-tests, unpaired for between subject and paired for within subject comparisons. Experiments with more than two groups that passed normality test were subjected to one-way ANOVA, two-way ANOVA or repeated measure two-way ANOVA, followed by Sidak’s tests for multiple comparisons when an interaction was found significant. Results that didn’t pass the normality test were analyzed using non parametric Friedman analysis, followed by Dunn’s multiple comparisons when an effect was found significant. Behavioral data for both initial contingency and reversed contingency tests were analyzed using a contrast analysis approach using PSY software (UNSW) as it allows for a more fine-grained data analysis of complex experimental designs. When an interaction was found, simple effect analysis was evaluated using PSY software. A value of p<0.05 was considered statistically significant. Graphs were originated using Graph Pad Prism 8. All other data were analyzed using Prism, Version 8.0 (Graph Pad software).

## Author contributions

Conceptualization: SB, BC, BWB

Methodology: SB, BC, LAB, RC, BKL

Investigation: SB, BC, LAB, RC, BKL

Visualization: SB, BC, BKL

Funding acquisition: BWB, LAB

Project administration: SB

Supervision: BWB

Writing – original draft: SB, BC

Writing – review & editing: BWB, LAB, BKL, RC

## Data and materials availability

All data are available in the supplementary materials.

## SUPPLEMENTARY MATERIALS

### Supplementary Figures S1 to S5

**Fig. S1.**
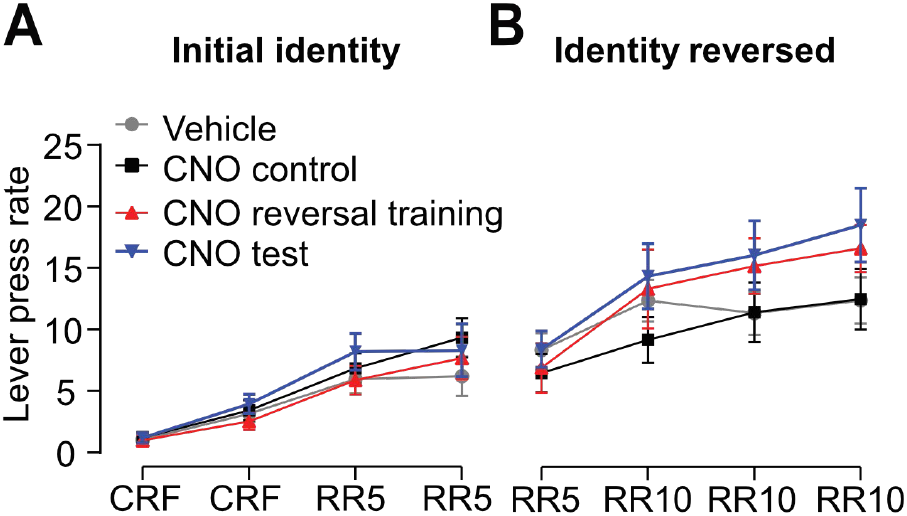
Effect of DREADD-induced inhibition of PL-pDMS pathway. Lever press training **(A)** prior to identity reversal, i.e., under the initial AO identity, and **(B)** after identity reversal (reversed identity) for each of the groups used in this experiment.

**Fig. S2.**
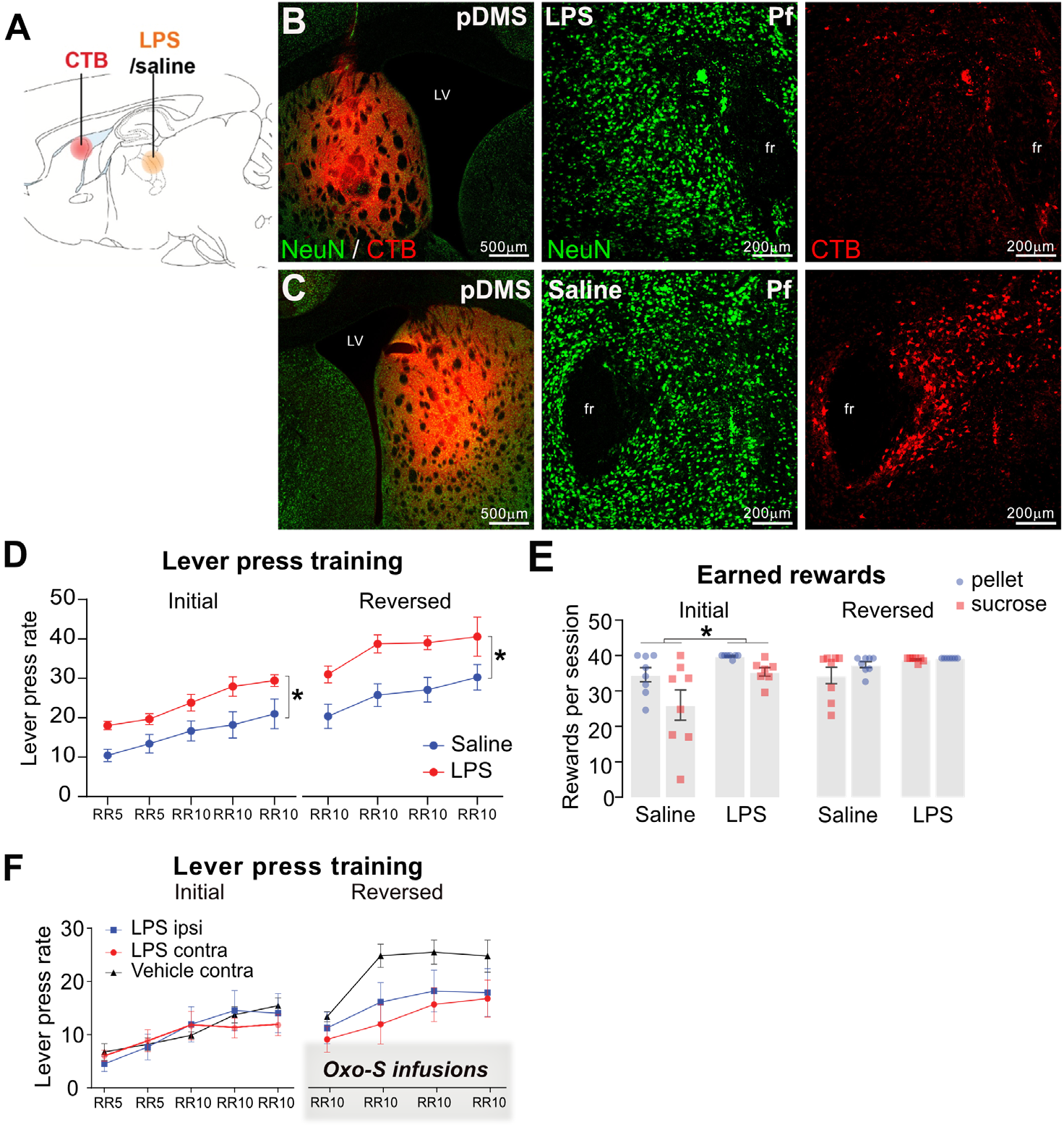
LPS effect on thalamostriatal pathway. **(A)** Illustration of the experimental design. Rats received CTB injected in the pDMS and LPS or saline injected in the Pf. (**B-C**) Representative micrograph of CTB injection in the pDMS (left panels) and saline or LPS in the Pf (middle and right panels). (**D**) Lever press rate during instrumental training after rats received a bilateral injection in the Pf of either saline or LPS. Rats quickly learned to press the levers and increased their performance as the ratio requirement increased. Statistical analysis showed an effect of training F(2.732,35.52)=17.23, p< 0.001 (Geisser-Greenhouse’s correction) and an effect of LPS F(1,13)=7.483, p=0.017 but no interaction between the two factors F<1. (**E**) Outcome earned during instrumental training of rats with saline or LPS injected bilaterally in the Pf. Statistically, we found an effect of LPS F(1,13)=5.314, p=0.038 and an overall preference for pellet over sucrose F(1,13)=12.57, p=0.004 during the initial AO identity. No differences were found during the identity reversal. All groups earned equal number of rewards during both initial and reversed AO identities. (**F**) Lever press rate during instrumental training. Rats received unilateral injections of Oxo-S in pDMS and LPS in the Pf injected ipsilaterally (LPS Ipsi) or contralaterally (LPS Contra) or they received Saline and Vehicle injected contralaterally (Saline Contra). All groups responded similarly during initial acquisition showing an effect of training F(2.684, 48.31)=17.18, p<0.001 and no main effect of group F(2,18)<1, nor interaction F(8, 72)=1.081, p>0.05. Training on the reversed identity revealed an overall effect of acquisition F(2.298, 41.36)=17.98 p<0.001 and no main effect of group F(2,18)=2.310, p>0.05, nor an interaction between training and group F(6,54)=1.392, p>0.05. The graph represents the mean of lever presses per minute ±1 SEM during initial AO identity training and during the reversed identity training. * indicates p<0.05.

**Fig. S3.**
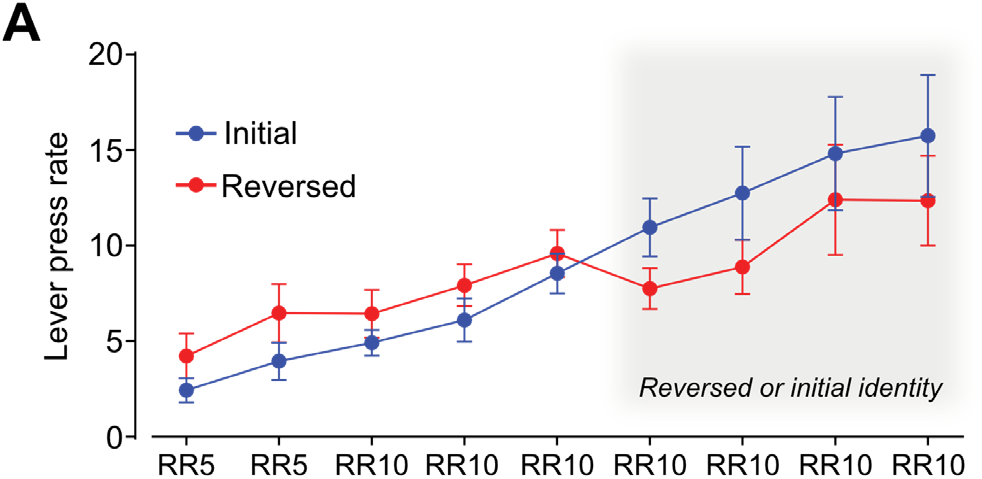
Training data. (**A**) Lever press rate during instrumental training in rats that were trained on the initial AO identity (Initial) and rats that were trained under initial identity followed by identity reversed (Reversed). We found an effect of training during both initial or reversed contingencies F(2.085, 37.52)=18.03, p<0.001 and F(2.162, 38.91)=12.90 p<0.001 respectively (Geisser-Greenhouse’s correction for both) and no training x group nor group differences between the two groups during both initial and reversed training session. Initial n=10, Reversed n=10. Error bars represent ±1 SEM. * indicates p<0.05.

**Fig. S4.**
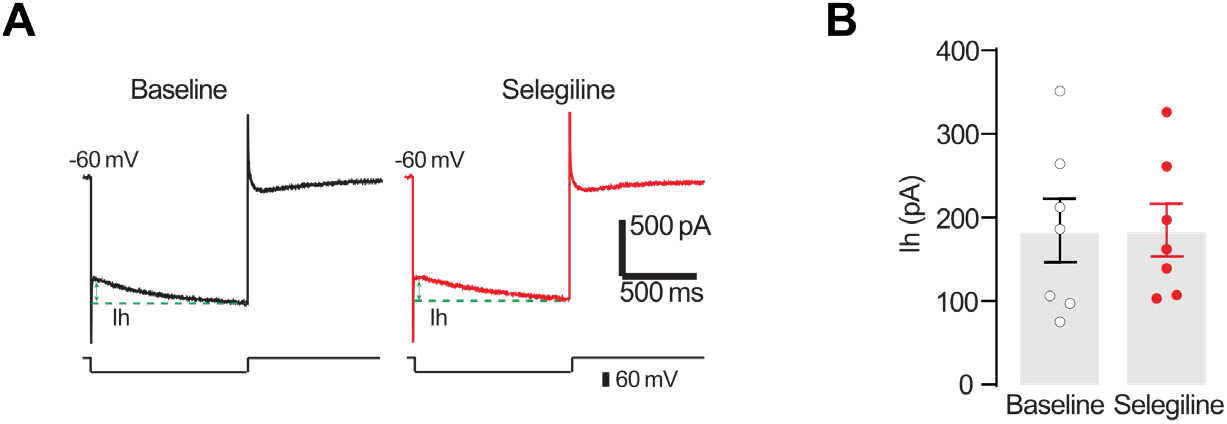
Selegiline has no effect on the hyperpolarization-activated cation current (Ih). CINs from naïve rats were held at −60mV under voltage-clamp in whole-cell configuration and a square-shaped hyperpolarizing voltage step of −60mV was injected into the cell via the recording pipette. (**A**) The raw current traces are direct result from corresponding voltage step applied (B). (**B**) Panel is the grouped data for baseline and during selegiline application. Two tailed paired *t*-test t=0.057, df=6, p>0.05, n=7. The graphs represent mean ±1 SEM.

**Fig. S5.**
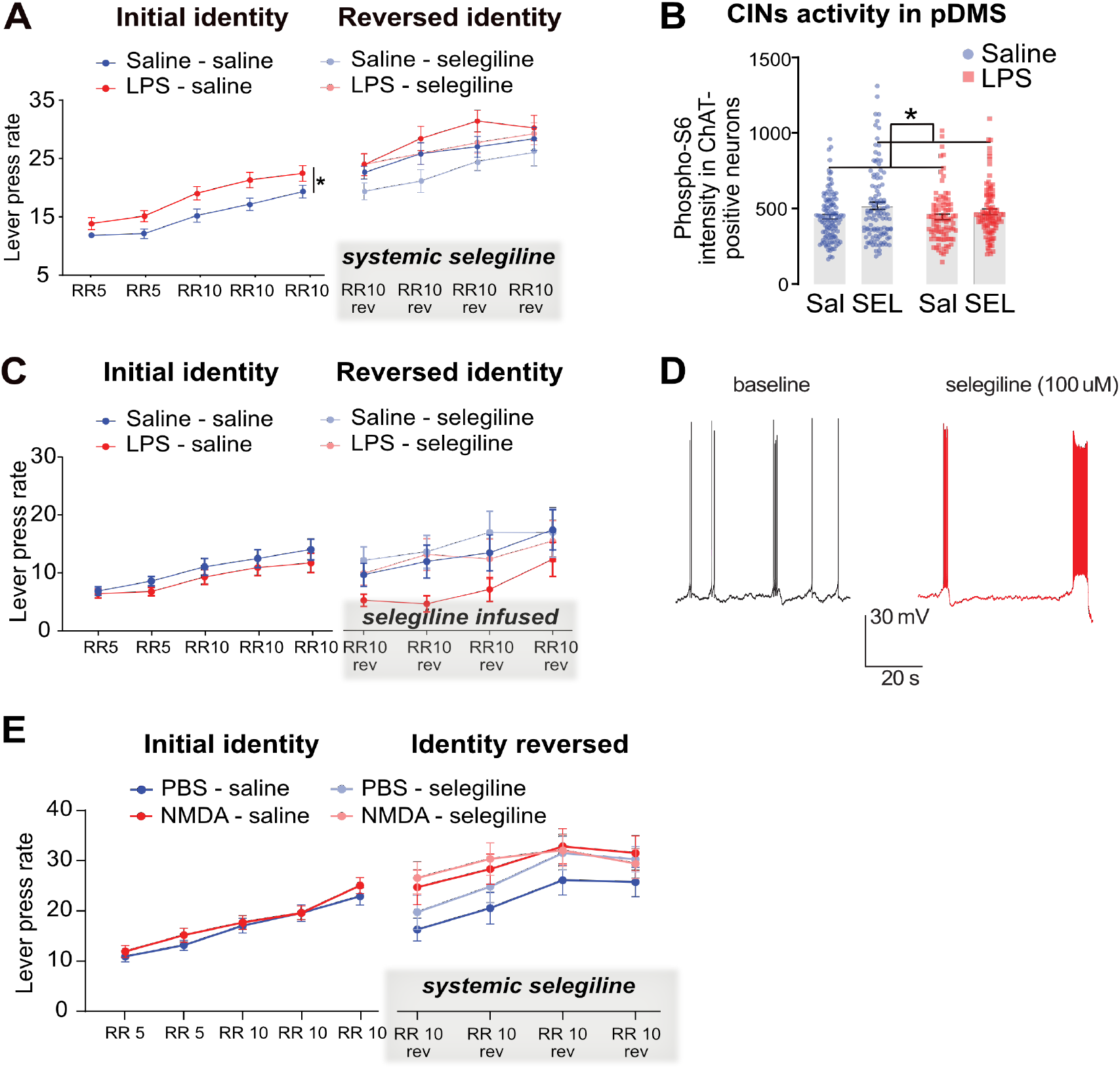
Selegiline rescues behavioral deficit in NMDA model of impaired flexible goal-directed control. (**A**) Lever press rate during instrumental training after rats received a bilateral injection in the Pf of either saline or LPS. Both saline and LPS groups showed a linear increase in pressing rate during sessions, increasing their performance as the ratio requirement increased. Statistical analysis revealed that LPS lesions increased lever pressing during initial acquisition F(1,46)=5.416 p=0.024; main effect of training F(3.013, 138.6)=76.80, p<0.001 but no group x training interaction F(4,184)=1.190, p>0.05. An effect of training was also observed during reversed training F(3, 126)=20, p<0.001, but no effect of LPS F(1,42)=3.699, p=0.06, nor an effect of selegiline F(1,42)=2.135, p>0.05, nor any interaction between these three variables, all Fs<1. **(B)** Quantification of p-S6rp in CINs after treatment with selegiline. Repeated measure ANOVA showed an effect of selegiline F(1, 408)=7.919, p<0.005, but no interaction LPS x selegiline F(1,408)=1.019, p>0.05. Saline Sal n=117 cells from 12 rats, Saline SEL n=109 cells from 10 rats, LPS Sal n=88 cells from 12 rats, LPS SEL n=98 cells from 10 rats. (**C**) Lever press rate during instrumental training after rats received a bilateral injection in the Pf of either saline or LPS and selegiline infusion in the pDMS. Both saline and LPS groups showed a linear increase in pressing rate during training, with an overall effect of training F(2.423,77.52)=22.75, p<0.001 (Geisser-Greenhouse’s correction) and no other significant effects. Rats were then trained on a reversed contingency, receiving infusion of selegiline 20 min before the start of each session. Statistical analysis revealed a clear training effect F(3, 90)=19.49 p<0.001, but no effect of LPS, nor of selegiline, nor interaction between these three variables, all Fs<1. **(D)** Selegiline-induced burst firing in NMDA model (n=4). **(E)** Lever press rate during instrumental training after rats received a bilateral injection in the Pf of either PBS or NMDA. Repeated measure ANOVA analysis of initial contingency showed a main effect of training F(2.940, 117.6)=84.14, p<0.001 (note: these statistics are Greenhouse-Geisser corrected for violations of sphericity), but no effect of lesion F(1, 40)=1.937, p>0.05, nor lesion x training interaction F(4, 160)=1.234, p>0.05. Statistical analysis of training on the reversed contingency revealed a main training effect F(3, 99)=39.47, p<0.001 no effect of NMDA lesion F(1, 34)<1, nor an effect of selegiline F(1, 33)=1.180, p>0.05, but an interaction between selegiline x training F(3, 99)=3.879, p=0.01 was found. The graph represents the mean of lever presses per minute ±1 SEM during initial AO identity training and during the reversed identity training. Selegiline was administered ip before each session of reversed identity training. Graphs represent average ±1 SEM. * indicates p<0.05.

## References and Notes

1. H. H. Yin, B. J. Knowlton, B. W. Balleine, Blockade of NMDA receptors in the dorsomedial striatum prevents action-outcome learning in instrumental conditioning. Eur J Neurosci 22, 505–512 (2005).

2. B. W. Balleine, A. Dickinson, Goal-directed instrumental action: contingency and incentive learning and their cortical substrates. Neuropharmacology 37, 407–419 (1998).

3. J. Peak, B. Chieng, G. Hart, B. W. Balleine, Striatal direct and indirect pathway neurons differentially control the encoding and updating of goal-directed learning. Elife 9, 1–28 (2020).

4. G. Hart, L. A. Bradfield, S. Y. Fok, B. Chieng, B. W. Balleine, The Bilateral Prefronto-striatal Pathway Is Necessary for Learning New Goal-Directed Actions. Current Biology 28, 2218–2229.e7 (2018).

5. Y. Hori, Y. Nagai, K. Mimura, T. Suhara, M. Higuchi, S. Bouret, T. Minamimoto, D1-and D2-like receptors differentially mediate the effects of dopaminergic transmission on cost benefit evaluation and motivation in monkeys. PLoS Biol 19 (2021), doi:10.1371/journal.pbio.3001055.

6. L. A. Bradfield, J. Bertran-Gonzalez, B. Chieng, B. W. Balleine, The Thalamostriatal Pathway and Cholinergic Control of Goal-Directed Action: Interlacing New with Existing Learning in the Striatum. Neuron 79, 153–166 (2013).

7. M. Matamales, Z. Skrbis, R. J. Hatch, B. W. Balleine, J. Götz, J. Bertran-Gonzalez, Aging-Related Dysfunction of Striatal Cholinergic Interneurons Produces Conflict in Action Selection. Neuron 90, 362–373 (2016).

8. S. R. Lapper, J. P. Bolam, Input from the frontal cortex and the parafascicular nucleus to cholinergic interneurons in the dorsal striatum of the rat (1992).

9. S. Consolo, P. Baronio, G. Guidit, G. di Chiara~, Role of the parafascicular thalamic nucleus and N-Methyl-D-Aspartate transmission in the D1-dependent control of in vivo acetylcholine release in rat striatum (1996).

10. S. Consolo, G. Baldi, S. Giorgi, L. Nannini, The Cerebral Cortex and Parafascicular Thalamic Nucleus Facilitate In Mvo Acetylcholine Release in the Rat Striatum through Distinct Glutamate Receptor Subtypes (1996).

11. J. P. Bolam, B. H. Wainer, A. D. Smith, Characterization of cholinergic neurons in the rat neostriatum. A combination of choline acetyltransferase immunocytochemistry, Golgi-impregnation and electron microscopy. Neuroscience 12, 711–718 (1984).

12. C. Contant, D. Umbriaco, S. Garcia, K. C. Watkins, L. Descarries, Ultrastructural characterization of the acetylcholine innervation in adult rat neostriatum. Neuroscience 71, 937–947 (1996).

13. J. B. Ding, J. N. Guzman, J. D. Peterson, J. A. Goldberg, D. J. Surmeier, Thalamic gating of corticostriatal signaling by cholinergic interneurons. Bone 23, 1–7 (2014).

14. B. W. Balleine, in Progress in Brain Research, (Elsevier B.V., 2022), vol. 269, pp. 227–255.

15. G. M. Halliday, Thalamic changes in Parkinson’s disease. Parkinsonism Relat Disord 15, S152–S155 (2009).

16. J. M. Henderson, S. B. Schleimer, H. Allbutt, V. Dabholkar, D. Abela, J. Jovic, M. Quinlivan, Behavioural effects of parafascicular thalamic lesions in an animal model of parkinsonism. Behavioural brain research 162, 222–232 (2005).

17. J. M. Henderson, K. Carpenter, H. Cartwright, G. M. Halliday, Degeneration of the centré median-parafascicular complex in Parkinson’s disease. Ann Neurol 47, 345–352 (2000).

18. D. Brooks, G. M. Halliday, Intralaminar nuclei of the thalamus in Lewy body diseases. Brain Res Bull 78, 97–104 (2009).

19. R. M. Villalba, T. Wichmann, Y. Smith, Neuronal loss in the caudal intralaminar thalamic nuclei in a primate model of Parkinson’s disease. Brain Struct Funct 219, 381–394 (2014).

20. M. A. Hely, W. G. J. Reid, M. A. Adena, G. M. Halliday, J. G. L. Morris, The Sydney multicenter study of Parkinson’s disease: The inevitability of dementia at 20 years. Movement Disorders 23, 837–844 (2008).

21. A. H. V. Schapira, K. R. Chaudhuri, P. Jenner, Non-motor features of Parkinson disease. Nat Rev Neurosci 18, 509–509 (2017).

22. D. Aarsland, K. Andersen, J. P. Larsen, A. Lolk, P. Kragh-Sørensen, Prevalence and characteristics of dementia in Parkinson disease: an 8-year prospective study. Arch Neurol 60, 387–92 (2003).

23. A. Surendranathan, J. B. Rowe, J. T. O’Brien, Neuroinflammation in Lewy body dementia. Parkinsonism Relat Disord 21, 1398–1406 (2015).

24. G. Gelders, V. Baekelandt, A. van der Perren, Linking neuroinflammation and neurodegeneration in parkinson’s diseaseJ Immunol Res 2018 (2018), doi:10.1155/2018/4784268.

25. T. Togo, E. Iseki, W. Marui, H. Akiyama, K. Uéda, K. Kosaka, Glial involvement in the degeneration process of Lewy body-bearing neurons and the degradation process of Lewy bodies in brains of dementia with Lewy bodies. J Neurol Sci 184, 71–75 (2001).

26. S. Iannaccone, C. Cerami, M. Alessio, V. Garibotto, A. Panzacchi, S. Olivieri, G. Gelsomino, R. M. Moresco, D. Perani, In vivo microglia activation in very early dementia with Lewy bodies, comparison with Parkinson’s disease. Parkinsonism Relat Disord 19, 47–52 (2013).

27. M. Deschênes, J. Bourassa, V. Doan, A. Parent, A single-cell study of the axonal projections arising from the posterior intralaminar thalamic nuclei in the rat. Eur J Neurosci 8, 329–343 (1996).

28. H. J. Groenewegen, H. W. Berendse, The specificity of the “nonspecific” midline and intralaminar thalamic nuclei. Trends Neurosci 17, 52–57 (1994).

29. G. Mandelbaum, J. Taranda, T. M. Haynes, D. R. Hochbaum, K. W. Huang, M. Hyun, K. Umadevi Venkataraju, C. Straub, W. Wang, K. Robertson, P. Osten, B. L. Sabatini, Distinct Cortical-Thalamic-Striatal Circuits through the Parafascicular Nucleus. Neuron 102, 636–652.e7 (2019).

30. B. N. Armbruster, X. Li, M. H. Pausch, S. Herlitze, B. L. Roth, Evolving the lock to fit the key to create a family of G protein-coupled receptors potently activated by an inert ligand (2007; www.pnas.orgcgidoi10.1073pnas.0700293104).

31. J. Bertran-Gonzalez, B. C. Chieng, V. Laurent, E. Valjent, B. W. Balleine, Striatal Cholinergic Interneurons Display Activity-Related Phosphorylation of Ribosomal Protein S6. PLoS One 7 (2012), doi:10.1371/journal.pone.0053195.

32. L. A. Bradfield, B. W. Balleine, Thalamic control of dorsomedial striatum regulates internal state to guide goal-directed action selection. Journal of Neuroscience 37, 3721–3733 (2017).

33. P. Calabresi, D. Centonze, A. Pisani, G. Sancesario, R. A. North, G. Bernardi, Muscarinic IPSPs in rat striatal cholinergic interneurones. J Physiol 510, 421 (1998).

34. V. Bernard, O. Laribi, A. I. Levey, B. Bloch, Subcellular Redistribution of m2 Muscarinic Acetylcholine Receptors in Striatal Interneurons In Vivo after Acute Cholinergic Stimulation. The Journal of Neuroscience 18, 10207 (1998).

35. M. E. Ragozzino, E. G. Mohler, M. Prior, C. A. Palencia, S. Rozman, Acetylcholine Activity in Selective Striatal Regions Supports Behavioral Flexibility. Neurobiol Learn Mem 91, 13 (2009).

36. Y. Smith, D. J. Surmeier, P. Redgrave, M. Kimura, Thalamic contributions to basal ganglia-related behavioral switching and reinforcement. Journal of Neuroscience 31, 16102–16106 (2011).

37. J. B. Ding, J. N. Guzman, J. D. Peterson, J. A. Goldberg, D. J. Surmeier, Thalamic gating of corticostriatal signaling by cholinergic interneurons. Neuron 67, 294–307 (2010).

38. S. Threlfell, T. Lalic, N. J. Platt, K. A. Jennings, K. Deisseroth, S. J. Cragg, Striatal Dopamine Release Is Triggered by Synchronized Activity in Cholinergic Interneurons (2012; http://www.sciencedirect.com/science/article/pii/S0896627312004436).

39. T. Aosaki, K. Kiuchi, Y. Kawaguchi, Dopamine D 1-Like Receptor Activation Excites Rat Striatal Large Aspiny Neurons In Vitro (1998).

40. P. Deng, Y. Zhang, Z. C. Xu, Involvement of Ih in dopamine modulation of tonic firing in striatal cholinergic interneurons. Journal of Neuroscience 27, 3148–3156 (2007).

41. P. Calabresi, B. Picconi, L. Parnetti, M. di Filippo, Personal View A convergent model for cognitive dysfunctions in Parkinson’s disease: the critical dopamine-acetylcholine synaptic balance (2006; http://neurology.thelancet.com).

42. J. A. Goldberg, J. N. J. Reynolds, Spontaneous firing and evoked pauses in the tonically active cholinergic interneurons of the striatumNeuroscience 198, 27–43 (2011).

43. B. D. Bennett, C. J. Wilson, Spontaneous Activity of Neostriatal Cholinergic Interneurons In Vitro. The Journal of Neuroscience 19, 5586 (1999).

44. K. Lee, A. K. Dixon, T. C. Freeman, P. J. Richardson, Identification of an ATP-sensitive potassium channel current in rat striatal cholinergic interneurones. J Physiol 510 (Pt 2), 441–453 (1998).

45. M. Beierlein, J. R. Gibson, B. W. Connors, Two Dynamically Distinct Inhibitory Networks in Layer 4 of the Neocortex. J Neurophysiol 90, 2987–3000 (2003).

46. Y. Kang, T. Notomi, M. Saito, W. Zhang, R. Shigemoto, Bidirectional Interactions between H-Channels and Na+-K + Pumps in Mesencephalic Trigeminal Neurons. Journal of Neuroscience 24, 3694–3702 (2004).

47. S. Becchi, A. Buson, B. W. Balleine, Inhibition of vascular adhesion protein 1 protects dopamine neurons from the effects of acute inflammation and restores habit learning in the striatum. J Neuroinflammation 18 (2021), doi:10.1186/s12974-021-02288-8.

48. N. Matsumoto, T. Minamimoto, A. M. Graybiel, M. Kimura, Neurons in the thalamic CM-Pf complex supply striatal neurons with information about behaviorally significant sensory events. J Neurophysiol 85, 960–976 (2001).

49. P. Apicella, Leading tonically active neurons of the striatum from reward detection to context recognition Trends Neurosci 30, 299–306 (2007).

50. C. Antoniades, H. Carageorgiou, S. Tsakiris, Effects of (-)deprenyl (selegiline) on acetylcholinesterase and Na(+),K(+)-ATPase activities in adult rat whole brain. Pharmacol Res 46, 165–169 (2002).

51. J. A. Goldberg, C. J. Wilson, Control of spontaneous firing patterns by the selective coupling of calcium currents to calcium-activated potassium currents in striatal cholinergic interneurons. Journal of Neuroscience 25, 10230–10238 (2005).

52. Y. Kawaguchi, Physiological, Morphological, and Histochemical Characterization of Three Classes of Interneurons in Rat Neostriatum (1993).

53. B. D. Bennett, J. C. Callaway, C. J. Wilson, Intrinsic Membrane Properties Underlying Spontaneous Tonic Firing in Neostriatal Cholinergic Interneurons (2000).

54. C. Kilkenny, W. Browne, I. C. Cuthill, M. Emerson, D. G. Altman, Animal research: Reporting in vivo experiments: The ARRIVE guidelines. Br J Pharmacol 160, 1577–1579 (2010).

55. J. C. McGrath, E. Lilley, Implementing guidelines on reporting research using animals (ARRIVE etc.): New requirements for publication in BJP. Br J Pharmacol 172, 3189–3193 (2015).

56. G. Paxinos, C. Watson, The rat brain in stereotaxic coordinates, 6th Edition | George Paxinos, Charles Watson | ISBN 9780080475158 (2006).

